# Control of 3’ Splice Site Selection in *S. cerevisiae* by a Highly Conserved Amino Acid within the Prp8 α-finger Domain

**DOI:** 10.1101/2025.06.16.659908

**Authors:** Ye Liu, Joshua C. Paulson, Aaron A. Hoskins

**Author notes:** **CORRESPONDING AUTHOR:** Aaron A. Hoskins.

## Abstract

Precise recognition of the boundaries between exons and introns (splice sites, SS) is essential for the fidelity of gene expression. In contrast with the 5’SS, the consensus 3’SS sequence in both *S. cerevisiae* and humans is just three nucleotides long: YAG. How the correct 3’SS is chosen among many possible alternates by the spliceosome is often unclear but likely involves proofreading by the Prp22 ATPase. In cryo-EM structures of spliceosome product (P) complexes, Glutamine 1594 in the highly conserved α-finger domain of the Prp8 protein interacts directly with the −3 pyrimidine of the 3’SS. To investigate the role of this interaction, we constructed a Prp8^Q1594A^ mutant and studied the impact on splicing and 3’SS selection. Using splicing reporter assays and RNA-seq, we show that Prp8^Q1594A^ enables use of non-consensus 3’SS by relaxing sequence requirements at the −3 and −2 positions. Consequently, this can change how adjacent 3’SS compete with one another during mRNA formation. The ability for Prp8^Q1594A^ to support splicing at non-YAG sites depends on the splicing factors Prp18 and Fyv6, and Prp8^Q1594A^ has genetic interactions with Prp22 mutants. Together, these findings suggest that the Prp8 α-finger acts as a sensor of 3’SS accommodation within the spliceosome active site. We propose that conformational change of the α-finger either allows or inhibits binding of the Prp22 c-terminal domain. This may provide a mechanism for regulating Prp22 activity in response to 3’SS binding.

## INTRODUCTION

The removal of introns and splicing of flanking exons is a critical step in gene expression in eukaryotes. This process occurs in two transesterification steps: 5’ splice site (5’SS) cleavage during which the phosphodiester bond between the 5’ exon and intron is broken concomitant with intron lariat formation and exon ligation during which a new phosphodiester bond is formed between the 5’ and 3’ exons and the intron lariat is released (**Fig. 1A**). Splicing is carried out by a large, highly conserved complex of small nuclear RNAs (snRNAs) and proteins called the spliceosome. It is essential for the fidelity of gene expression that splicing be carried out at precisely defined locations—a single nucleotide (nt) error in choice of the 5’ or 3’SS can destroy the reading frame of a mRNA and prevent protein production. As a result, the splicing machinery uses multiple recognition and proofreading strategies to ensure that 5’ and 3’SS, as well as the site of lariat formation (the branch point sequence, BPS), are chosen correctly (Semlow and Staley 2012).

**Figure 1.**
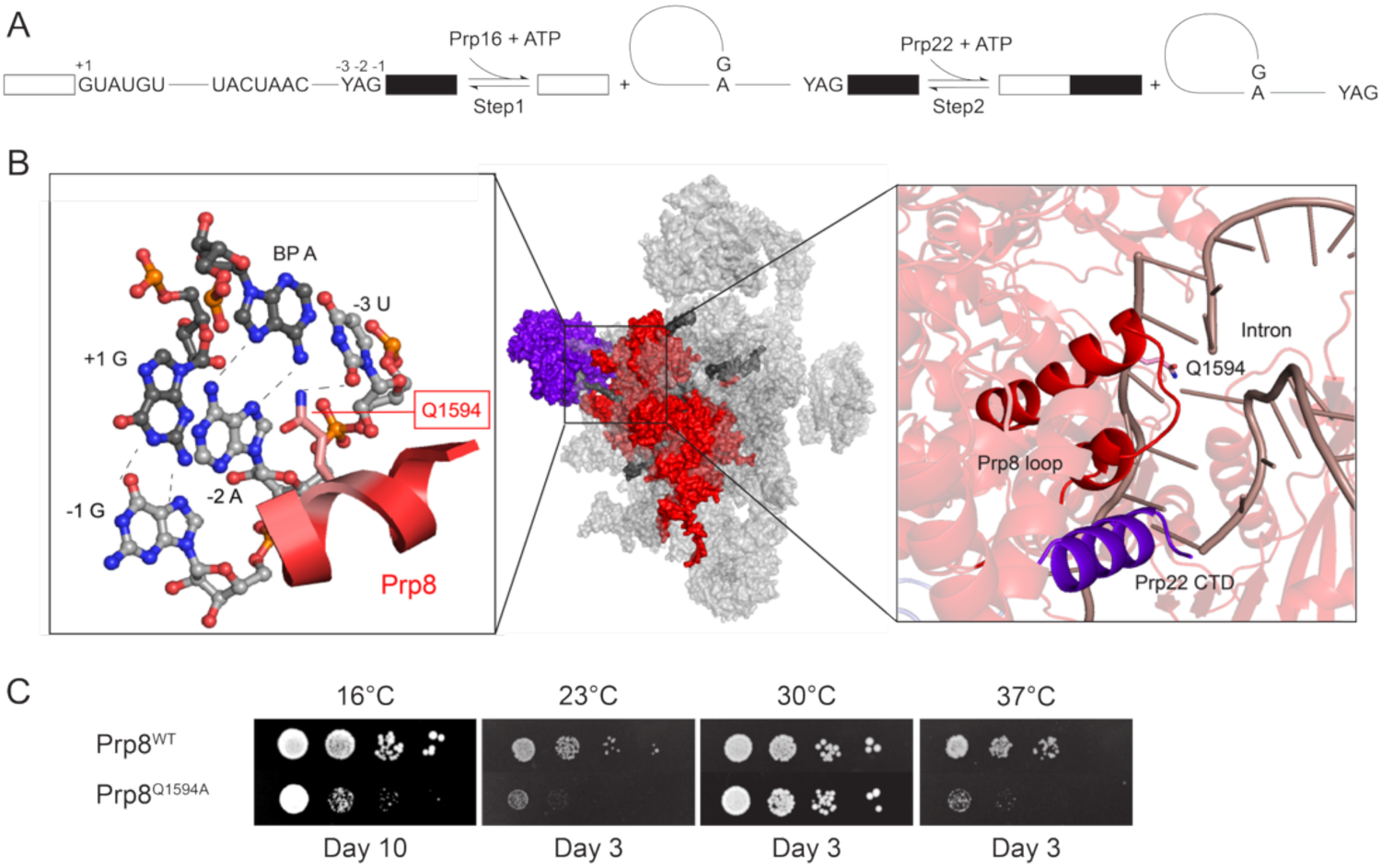
Prp8 Residue Q1594 Directly Contacts the 3’ SS. **(A)** Schematic diagram showing the two transesterification steps during pre-mRNA splicing. The 3’SS is numbered (−3 to −1) relative to the intron/exon boundary. **(B)** Structural basis of 3’SS recognition in the yeast P complex spliceosome (9DTR). Left panel: detailed interactions between the 3’SS UAG sequence and the intron (−1G and −2A positions) and Prp8^Q1594^ (- 3U position). Central panel: overall structure of the P complex spliceosome highlighting Prp8 (red), Prp22 (purple), and the spliced products and lariat intron (dark grey). Right panel: a close-up view of the interaction between the Prp8 α-finger domain loop, containing Q1594 (red), with the C-terminal domain (CTD) of Prp22 (purple). Figure generated with PyMOL (Schrödinger, LLC). **(C)** Growth assays comparing wild-type (WT) Prp8 and the Q1594A mutant at different temperatures. YPD plates were incubated at the given temperatures and images collected on the noted days.

Relative to the 5’SS and BPS, recognition of the 3’SS is quite challenging for the splicing machinery. Consensus 3’SS motifs are short, just three nucleotides: YAG. Further, YAG sequences can be common in introns meaning that the spliceosome must not only select a YAG sequence as a 3’SS but also choose the correct YAG among many possible alternates. A number of disease-associated mutations can lead to creation of aberrant 3’SS given the short consensus (Vorechovsky 2006). The complexity of this problem is best illustrated by the common NAGNAG 3’SS motif found in humans (Busch and Hertel 2012). This sequence contains two adjacent 3’SS; however, only one of which can be used to generate the correct mRNA isoform. NAGNAG sites occur in thousands of human 3’SS and represent the second most common form of alternative splicing in which the reading frame is conserved—often regulating the addition or removal of a single amino acid from a protein isoform (Bradley et al. 2012). For example, the loss of a single amino acid in patients with the 2623G>T mutation in the *CFTR* gene (to create a CAGuAG 3’SS) can lead to mild cystic fibrosis phenotypes and a 2588G>C mutation in the *ABCR* gene (to create a UAGcAG 3’SS) results in loss of a single amino acid in some patients with Stargardt disease, a type of macular dystrophy (Maugeri et al. 1999; Hinzpeter et al. 2010).

Cryo-EM of spliceosomes has revealed the molecular mechanism for 3’SS recognition (**Fig. 1B**) (Bai et al. 2017; Liu et al. 2017; Wilkinson et al. 2017). Unlike recognition of the 5’SS and BPS which are identified by base pairing to the U1 and U6 snRNAs (5’SS) or U2 snRNA (BPS), the 3’SS is recognized by non-Watson-Crick-Franklin base pairing. The terminal guanine of the 3’SS is recognized by Hoogsteen pairing to the first guanine of the intron (the 5’ terminal guanine of the 5’SS). The conserved adenine of the YAG motif is also recognized by Hoogsteen pairing but to the branch point adenosine of the intron. Finally, the pyrimidine nucleotide of the YAG motif is proposed to be recognized by a highly conserved glutamine (Q1594) located in the α-finger domain of the splicing factor Prp8. It has been proposed that preference for a pyrimidine at this position is largely due to sterics with larger purine nucleotides leading to clashes between the protein side chain and the 3’SS (Wilkinson et al. 2017). Despite this pattern of recognition, it is well-known that the splicing machinery can utilize other 3’SS sequences, especially in the presence of mutated protein or RNA splicing factors (Mayas et al. 2006; Liu et al. 2007; Query and Konarska 2012; Roy et al. 2023). This suggests that the spliceosome active site is flexible enough to accommodate a variety of interactions that result in juxtaposition of the 5’ and 3’SS.

When multiple 3’SS are present, it is believed that spliceosomes utilize a scanning mechanism to choose the site of exon ligation (Patterson and Guthrie 1991; Smith et al. 1993; Perez-Valle and Vilardell 2012; Semlow et al. 2016). In this mechanism, spliceosomes tend to select 3’SS based on their proximity to the BPS by scanning in a 5’ to 3’ direction as opposed to a “diffusion-collision” model in which the 3’SS would randomly dock into the active site (Umen and Guthrie 1995). Simple scanning cannot account for all observations of 3’SS usage *in vivo*, and proofreading by the Prp22 ATPase can result in “leaky scanning” due to ATP-dependent rejection of inefficient 3’SS (Semlow et al. 2016). During the rejection step, it is expected that the 3’SS is dissociated from the active site (*i.e.,* docking of substates for exon ligation is reversible (Abelson et al. 2010)) to allow other, competing 3’SS to bind. Interestingly, the c-terminal domain (CTD) of Prp22 is inserted into the core of the spliceosome and is located next to the Prp8 α-finger domain (**Fig. 1B**). While the function of the Prp22 CTD has not yet been rigorously studied, this proximity suggests that it plays a role in communicating docking of the 3’SS into the spliceosome active site and/or completion of exon ligation to Prp22.

Even with the leaky scanning mechanism, it is difficult to understand how alternative and adjacent 3’SS are selectively utilized by the splicing machinery in a tissue-specific manner (Hiller and Platzer 2008; Bradley et al. 2012). It has been proposed that in many cases RNA secondary structures or intronic sequence elements play roles in selection of one NAG site over another (Smith et al. 1993; Meyer et al. 2011). Recent cryo-EM, molecular biology, and RNA-seq experiments have also shown that a number of 2^nd^-step protein factors (proteins specifically recruited just prior to and/or playing a role in exon ligation) influence 3’SS choice during the catalytic stages of splicing in humans (Dybkov et al. 2023). While some of these factors (like Prp8, Prp22, Prp18, and Fyv6/FAM192A) are conserved between *Saccharomyces cerevisiae* (yeast) and humans (Roy et al. 2023; Senn et al. 2024), others are believed to be metazoan-specific including CACTIN, FAM50A, NKAP, PRKRIP1, and FAM32A. In comparison with the human splicing machinery, how yeast spliceosomes discriminate between competing NAGNAG sites is not well-studied.

Here, we have generated yeast strains containing a Prp8 Q1594A mutation (Prp8^Q1594A^) and studied the effects on pre-mRNA splicing. As predicted based on spliceosome cryo-EM structures, this mutation results in a loss of selectivity for pyrimidines at the –3 position of the 3’SS. Surprisingly, loss of selectivity is also seen at the –2 position indicating that the Prp8 α- finger domain regulates multiple aspects of SS usage. As a result of this mutation, usage of competing 3’SS can change due to increased utilization of non-YAG 3’SS. Accommodation of these sites in Prp8^Q1594A^-containing spliceosomes may be due to increased flexibility of the active site. Prp8^Q1594A^ mutant strains also exhibit genetic interactions with Prp22 mutations suggesting that Prp8 α-finger interactions with the 3’SS are important for regulating Prp22 activity. Finally, transcriptome analysis shows that the Prp8^Q1594A^ mutation results in increased usage of cryptic, non-YAG 3’SS across the spliced RNA transcriptome. Together our results provide direct evidence for function of the Prp8 α-finger in 3’SS selection and allow us to propose a model in which conformational change of this domain regulates splicing.

## RESULTS

### The Prp8^Q1594A^ Mutation Results in Temperature-Dependent Growth Defects

The Prp8 α-finger, including Q1594, is highly conserved (**Sup. Fig. S1**) consistent with a critical function during splicing. A prior study reported that a Prp8^Q1594A^ mutation resulted in normal growth at 30°C but did not include any other analysis of this mutant (Wilkinson et al. 2017). To study the role of this residue further, we first attempted to replicate this result as well as prepare additional Prp8^Q1594^ mutants including those that removed the side chain (Prp8^Q1594G^), reduced the side chain size but preserved its hydrogen bonding characteristics (Prp8^Q1594N^), or introduced a charged side chain at this position (Prp8^Q1594E^, Prp8^Q1594R^). We prepared TRP/CEN plasmids containing PRP8 genes with the corresponding mutations, transformed these into yeast in which the chromosomal copy of PRP8 was deleted and Prp8 expressed from a URA/CEN plasmid, and selected for growth on plates containing 5-FOA. We were only able to obtain colonies containing either Prp8^WT^ or Prp8^Q1594A^. For the other Prp8 mutants, we observed fewer colonies forming on the 5-FOA-containing plates, and DNA sequencing results showed that these yeasts contained Prp8^WT^ but not the desired mutations (data not shown). In agreement with the previous study, yeast containing Prp8^Q1594A^ grew similarly to Prp8^WT^ on YPD plates at 30°C (**Fig. 1C**). However, this mutation resulted in significant growth defects at higher (37°C; temperature sensitive or *ts*) and lower temperatures (16 and 23°C; cold sensitive or *cs*). These results indicate that while yeast containing Prp8^Q1594A^ are viable, this amino acid is important for yeast growth under certain conditions and that some amino acid substitutions at this position are likely not tolerated.

### Prp8^Q1594A^ Improves Splicing at Nonconsensus 3’ Splice Sites

To assess the impact of Prp8^Q1594A^ on splicing, we used the ACT1-CUP1 reporter system in which yeast tolerance to Cu^2+^ is proportional to splicing of a reporter pre-mRNA (**Fig. 2A**). We compared growth of strains containing either Prp8^WT^ or Prp8^Q1594A^ on plates containing increasing concentrations of Cu^2+^ (0–2.5mM) using reporters with either a consensus 3’SS (UAG) or substitutions at the −3, −2, or −1 positions (**Fig. 2B**, **C**). Yeast with Prp8^Q1594A^ showed lower Cu^2+^ tolerance relative to WT for the consensus reporter. This is consistent with a negative impact on splicing and decreased amounts of mRNA production.

**Figure 2.**
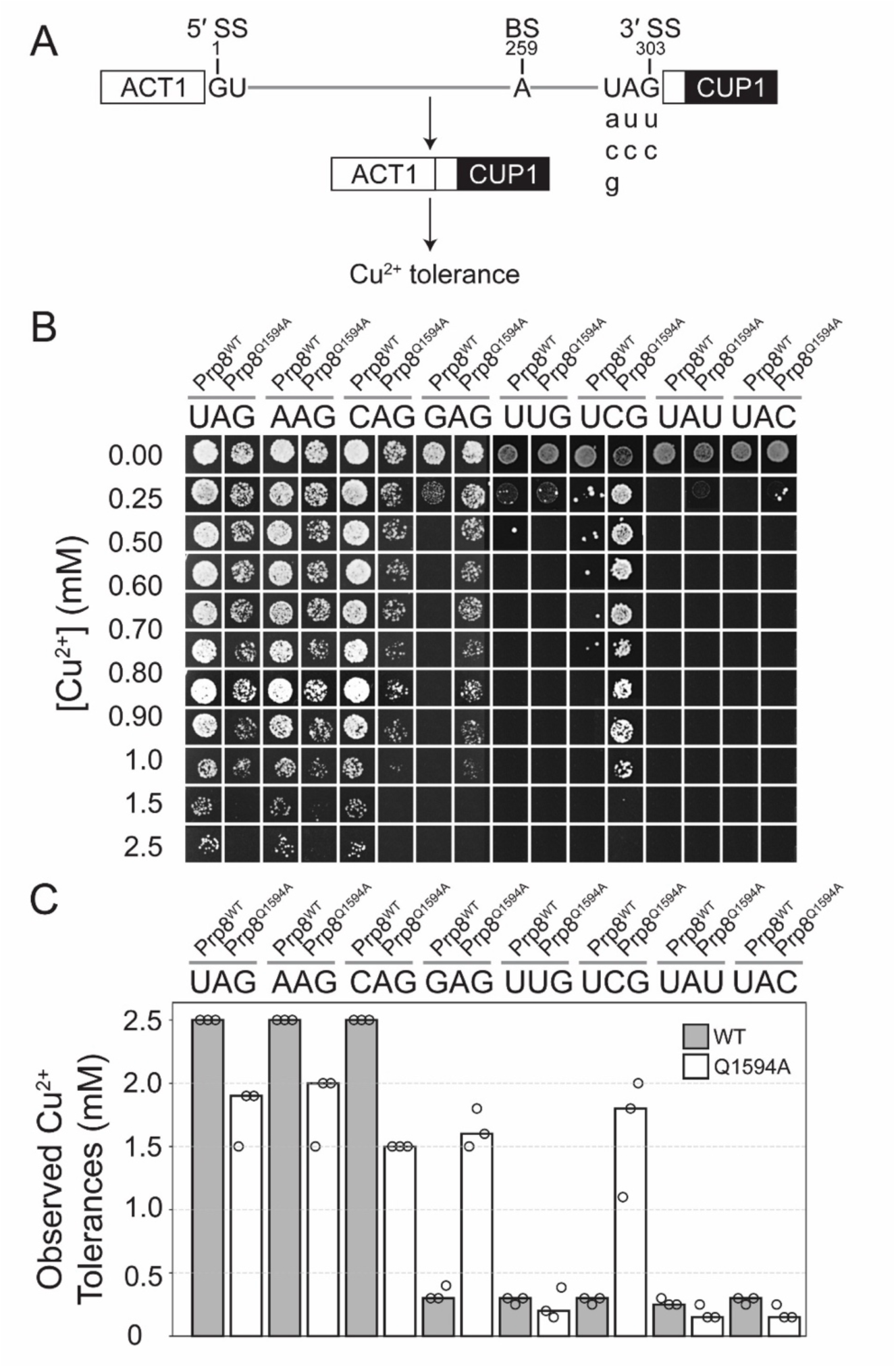
Prp8^Q1594A^ Increases Copper Tolerance for Reporters with Nonconsensus 3’ SS. **(A)** Schematic of the ACT1-CUP1 splicing reporter assay, with tested variants at the −1, −2, and −3 positions of the 3’ SS noted. **(B)** Representative growth of yeast strains expressing either Prp8^WT^ or the Q1594A mutant on plates containing varying concentrations of Cu²⁺ after 3 days at 30°C. **(C)** Observed Cu²⁺ tolerances for Prp8 strains containing the noted ACT-1CUP1 3’ SS reporters. Bar heights represent the values from a single biological replicate. Circles represent values obtained from three biological replicates.

Despite the negative impact on a consensus substrate, the Prp8^Q1594A^ mutation strongly improved Cu^2+^ tolerance and production of mRNA in yeast containing nonconsensus GAG and UCG 3’SS (**Fig 2B**, **C**; **Sup. Fig. S2**). Improvement of splicing of the reporter containing the GAG 3’SS is consistent with the proposal that Prp8^Q1594^ provides a steric block to accommodating a guanosine at the −3 position (Wilkinson et al. 2017). In this case, the introduction of a smaller, alanine side chain allows the guanosine to be more easily accommodated. Surprisingly, Prp8^Q1594A^also improved growth of a reporter containing a cytidine at the −2 position (UCG). It has been proposed that recognition of the −2 position in the consensus 3’SS is due to hydrogen bonding between the Hoogsteen edges of the −2 adenosine and branch point adenosine. We do not know if there is any interaction between the branch point adenosine and the −2 cytidine in the UCG reporter; however, A:C pairs can form with one or more hydrogen bonds, the latter requiring ionization or tautomerization of the nucleotides (Lemieux and Major 2002). The presence of a smaller residue at Prp8 position 1594 may allow additional flexibility within the spliceosome active site to accommodate these interactions or lack of interaction altogether. In contrast, we did not observe any increase in Cu^2+^ tolerance or mRNA production for the UUG reporter. This would indicate that any additional flexibility afforded by Prp8^Q1594A^ to facilitate splicing does not extend to uridines at this position.

Finally, we noted that strains containing either Prp8^WT^ or Prp8^Q1594A^ had very low copper tolerances in the presence of reporters with substitutions at the −1 site: UAU and UAC (**Fig. 2B, C**). However, the impacts on splicing of these substitutions are quite different between the strains. In the case of Prp8^WT^-containing strains, we observed significant mRNA production with both the UAU and UAC reporters despite the low copper tolerances (**Sup. Fig. S2**). This is likely due to activation of a cryptic, YAG-type 3’SS by introduction of these substitutions [UAG/AG becomes UA<u>U/AG</u> (new cryptic site underlined); similar for the UAC substitution]. The new 3’SS produces a change in reading frame of the mRNA and results in mRNA production but low levels of CUP1 protein production and Cu^2+^ tolerance. When Prp8^Q1594A^ was present, we observed little evidence for mRNA production with the UAU and UAC reporters. This would indicate that Prp8^Q1594A^ prevents recognition of the cryptic 3’SS in this context even though it has a consensus sequence.

### Prp8^Q1594A^ Changes Competition Between Potential Splice Sites

Given the unusual results with recognition of cryptic YAG-type 3’SS when they overlap with UAU or UAC sites, we next tested how the Prp8^Q1594A^ mutant would impact competition between two adjacent, NAGNAG-type 3’SS sites. We used the ACT1-CUP1 reporter assay and primer extension analysis to monitor 3’SS usage in reporters with two consensus sites (UAGUAG) or those in which a nonconsensus site is located proximal (GAGUAG) or distal to the BPS (UAGGAG; **Fig. 3A**).

**Figure 3.**
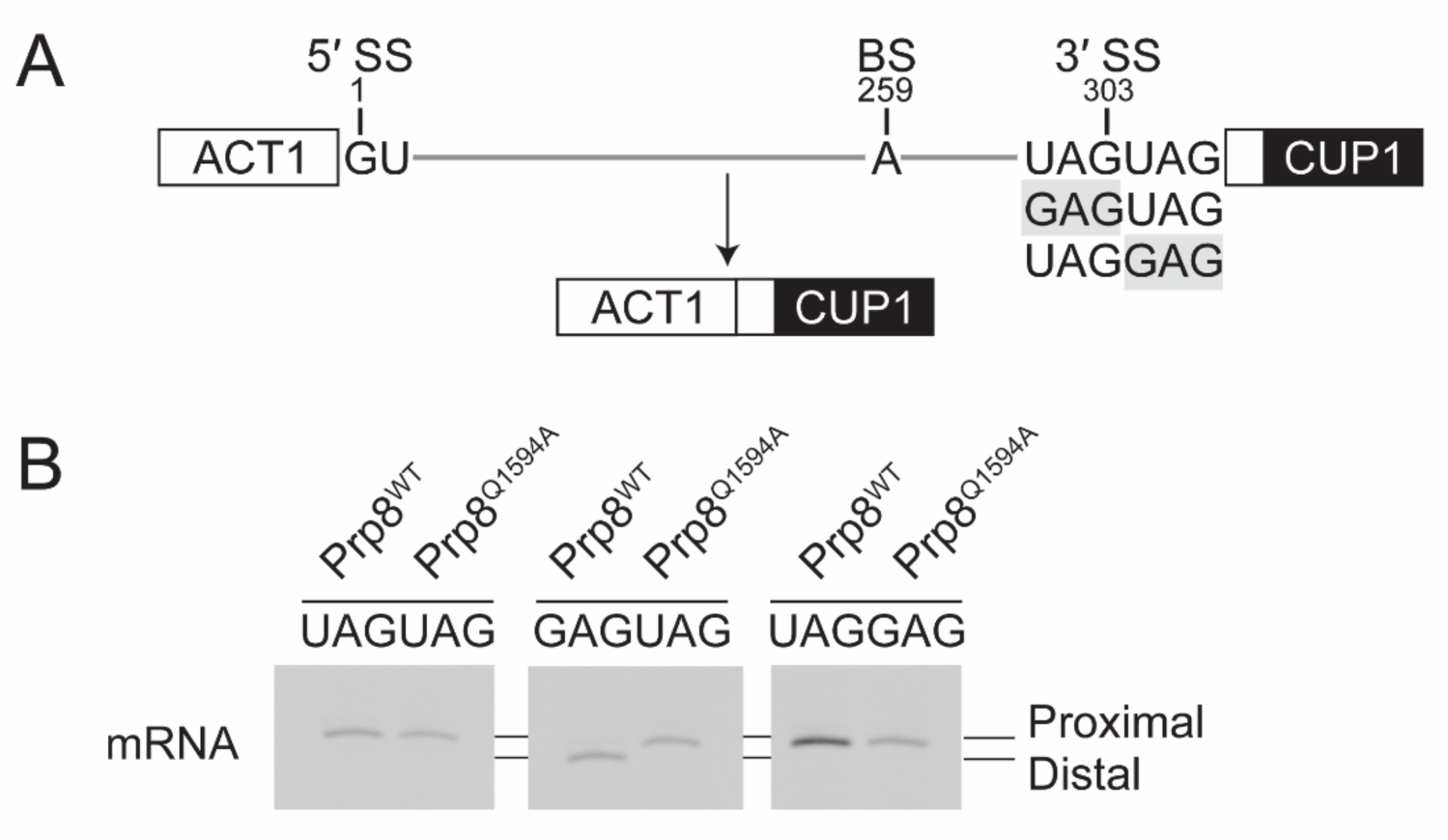
Prp8^Q1594A^ Alters Competition between Tandem 3’SS. **(A)** Schematic of the ACT1-CUP1 reporter construct with three tandem 3’SS variants The nonconsensus GAG site is highlighted in grey. **(B)** Representative primer extension assay results. Note that yeast with Prp8^WT^ utilize the distal UAG site, while yeast with Prp8^Q1594A^ use the proximal GAG site in the GAGUAG reporter. The images shown in this panel were all from the same gel but intervening lanes removed and lanes reordered for clarity.

As expected, strains containing either Prp8^WT^ or Prp8^Q1594A^ both showed exclusive use of the proximal UAG site with the UAGUAG reporter (**Fig. 3B**; **Sup. Fig. S3**). When a nonconsensus GAG site was included in the BPS proximal position (GAGUAG), the strain with Prp8^WT^ now made use of the distal, consensus UAG site while the strain with Prp8^Q1594A^ maintained used of the proximal (GAG) site. When the GAG site was moved to the distal position (UAGGAG), both strains showed preferred use of the BPS proximal, consensus site. Together, these results are consistent with a “scanning” model of 3’SS selection in which sites are primarily used based on their position relative to the BPS in a 5’ to 3’ direction. In the case of Prp8^WT^ strains, the GAG site in the GAGUAG reporter is either not recognized as a 3’SS or is recognized and then rejected in favor of the adjacent UAG site (likely by a Prp22-dependent mechanism) (Mayas et al. 2006). In the case of the Prp8^Q1594A^, results with the UAGGAG reporter indicate that a consensus UAG 3’SS is still recognized as “optimal” in this context and not subject to rejection in a manner that would facilitate use of the adjacent GAG site.

### Fyv6 and Prp18 Influence Use of Nonconsensus 3’ Splice Sites with Prp8^Q1594A^

Recognition of the 3’SS can also depend on 2^nd^-step factors such as the nonessential proteins Fyv6 and Prp18 (Lipinski et al. 2023; Roy et al. 2023). We next tested if changes in Fyv6 or Prp18 would exacerbate or suppress changes in Cu^2+^ tolerance due to Prp8^Q1594A^. We were able to create yeast strains containing Prp8^Q1594A^ but without Fyv6 (*fyv6Δ*) showing that this combination is not synthetic lethal. Loss of Fyv6 results in *cs* and *ts* growth defects, and these were not changed by Prp8^Q1594A^ (**Sup. Fig. S4**) (Lipinski et al. 2023; Roy et al. 2023). When analyzed using the ACT1-CUP1 assay, yeast containing Prp8^Q1594A^ but lacking Fyv6 failed to improve Cu^2+^ tolerance in the presence of the GAG or UCG 3’SS reporters (**Fig. 4A**, **B**; *cf.* **Fig. 2B**, **C**).

**Figure 4.**
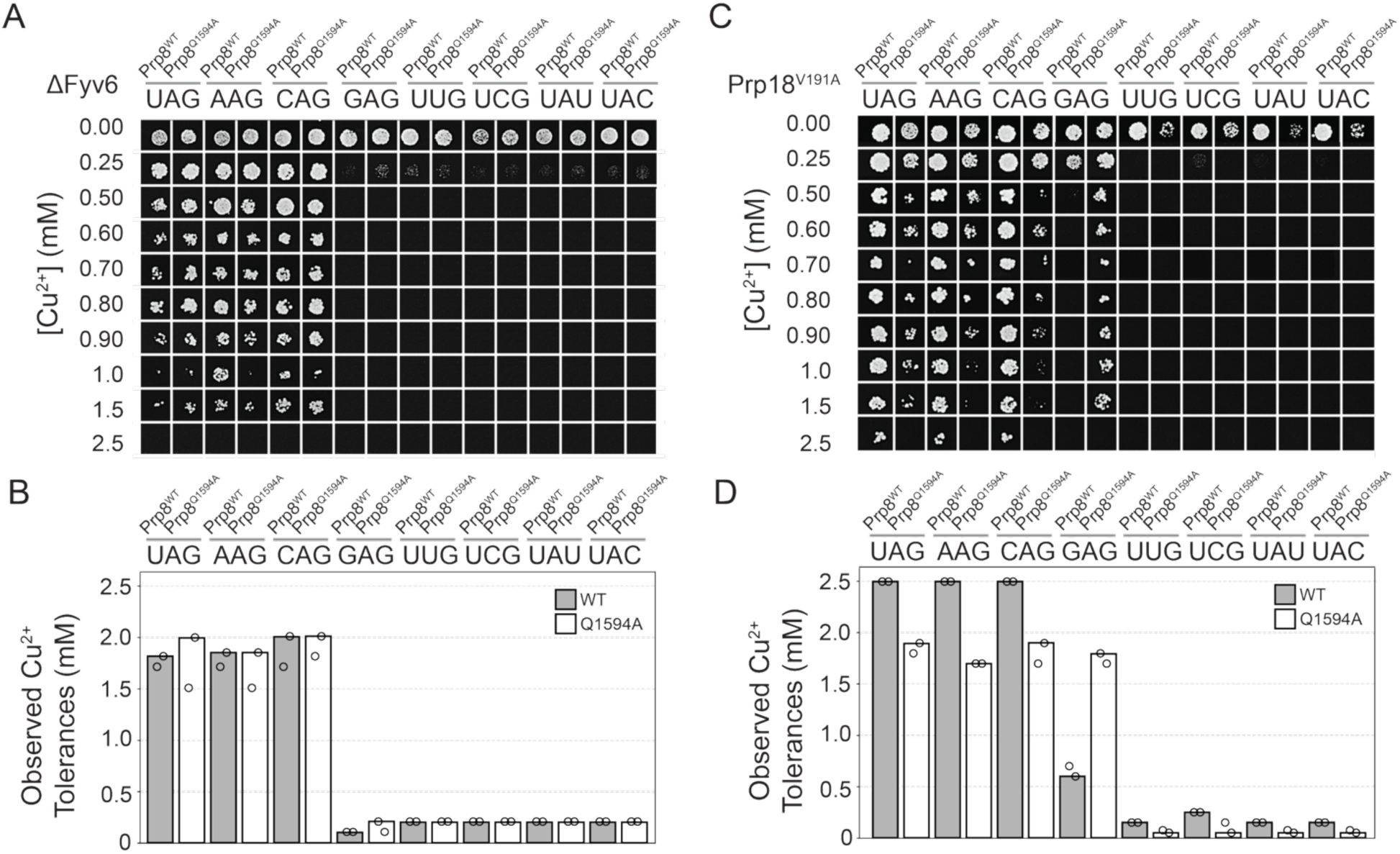
Fyv6 and Prp18 Impact Copper Tolerances of Yeast with Prp8^Q1594A^. **(A, C)** Representative growth of yeast strains in the ΔFyv6 (panel A) or Prp18^V191A^ (panel B) background expressing either WT Prp8 or the Q1594A mutant in varying copper ion concentrations ([Cu²⁺]). **(B, D)** Observed Cu²⁺ tolerances containing the noted ACT- 1CUP1 3’SS reporters. Bar heights represent the values from a single biological replicate. Circles represent values obtained from two biological replicates.

We were able to make similar observations using the dominant negative Prp18^V191A^ mutant (Aronova et al. 2007). The presence of this mutant Prp18 protein was also not lethal in combination with Prp8^Q1594A^. It did not change the *cs* phenotype due to Prp8^Q1594A^, but it did slightly suppress the *ts* phenotype at 37°C (**Sup. Fig. S4**). As with strains lacking Fyv6, we did not observe improved Cu^2+^ tolerance for the UCG 3’SS reporters in strains containing both Prp8^Q1594A^ and Prp18^V191A^; however, tolerance was still improved for the GAG reporter (**Fig. 4C**, **D**). Together these results show that Prp18 and Fyv6 influence nonconsensus 3’SS usage by Prp8^Q1594A^- containing spliceosomes. It is possible that accommodation of nonconsensus 3’SS within the spliceosome requires active site flexibility. This may be able to be achieved by alteration of Prp8 or 2^nd^-step factors. However, there might be a limit to how much flexibility the active site can tolerate, and the presence of multiple perturbations results in a failure to support some types of nonconsensus splicing.

### Genetic Interactions between Prp8^Q1594A^ and Prp22

Given the proximity of the Prp8 α-finger to the Prp22 CTD and the influence of Prp22 on 3’SS selection and proofreading (Mayas et al. 2006; Semlow et al. 2016; Roy et al. 2023), we tested if there would be genetic interactions between Prp8^Q1594A^ and Prp22 mutations. We chose three different mutations in Prp22 that have previously been characterized. Prp22^S635A^ is a mutation in DExD/H-box motif III that results in a *cs* phenotype in yeast (Schwer and Meszaros 2000). In assays with purified protein, Prp22^S635A^ enzyme shows near WT levels of ATP hydrolysis activity but is severely defective at unwinding RNA duplexes (Schwer and Meszaros 2000). This indicates the mutation likely disrupts coupling between ATP hydrolysis and RNA binding or unwinding. This mutation also permits splicing at nonconsensus 3’SS (Mayas et al. 2006). The G810A mutation in motif VI is also *cs,* likely impairs ATP hydrolysis, and is mRNA release defective (Schwer and Meszaros 2000; Mayas et al. 2006). It also permits usage of nonconsensus 3’SS *in vivo* including those with a gAG sequence (Mayas et al. 2006; Roy et al. 2023). Finally, the I1133R mutation is located in the CTD and partially suppresses *cs* and *ts* phenotypes due to loss of the 2^nd^-step factor Fyv6 (Senn et al. 2024).

We shuffled each of plasmids carrying genes for WT or mutant Prp22 proteins into yeast strains with either Prp8^WT^ or Prp8^Q1594A^. We then selected for growth on plates containing 5-FOA at 30°C. We were able to observe growth for both Prp8^WT^ or Prp8^Q1594A^-containing strains for Prp22^WT^, Prp22^G810A^, and Prp22^I1133R^. However, we observed no growth for a strain containing both Prp8^Q1594A^ and Prp22^S635A^ on FOA-containing plates at 16, 23, 30, or 37°C (data not shown). This indicates that there is a likely synthetic lethal genetic interaction between this Prp8 α-finger mutation and Prp22^S635A^.

Yeast containing Prp8^Q1594A^ and Prp22^G810A^ are *ts* and have a severe growth defect at 37°C (**Fig. 5**). The *ts* phenotype is strongly suppressed by the Prp22^I11133R^ CTD mutation. Together, these results indicate that genetic interactions can occur between mutations in the Prp8 α-finger and Prp22 active site over a distance of ∼ 90Å (Senn et al. 2024). It is possible that the origin of the phenotype could be due to production of misspliced mRNAs (Mayas et al. 2006; Roy et al. 2023) and/or failure to properly couple Prp22 activity with 3’SS or spliced product recognition by Prp8. The mechanism of suppression by Prp22^I11133R^ could involve changing how the CTD interacts with the spliceosome core (*e.g.,* stronger or weaker tethering of Prp22 to the spliceosome) and/or how this domain may be involved in regulating Prp22 activity (see **Discussion**).

**Figure 5.**
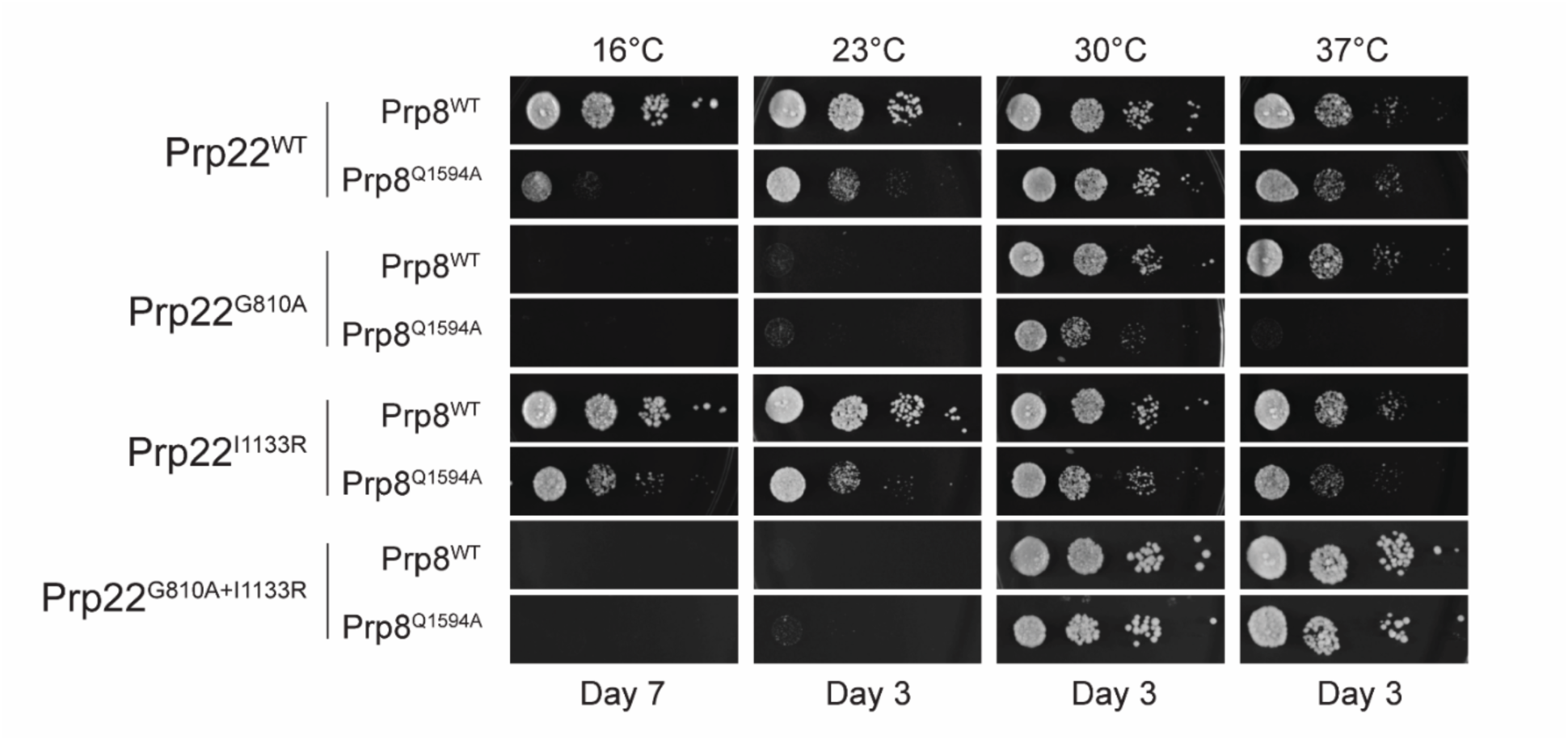
Genetic Interactions between Prp8^Q1594A^ and Prp22 Mutants. Growth assays comparing Prp8^WT^ and the Q1594A mutant at different temperatures in the presence of different Prp22 mutants. YPD plates were incubated at the given temperatures and images collected on the noted days.

### Prp8^Q1594^ is Critical for Proper 3’ Splice Site Usage Across the Yeast Transcriptome

To investigate the role of Prp8^Q1594A^ in transcriptome-wide 3’SS selection, we performed RNA-seq analysis comparing Prp8^WT^ and Prp8^Q1594A^ yeast strains, while knocking out UPF1 (*upf1Δ*) to reduce nonsense-mediated mRNA decay (NMD) (Kawashima et al. 2014). This knockout reduces degradation of aberrantly spliced mRNA (such as those with use of alternate 3’SS), allowing detection of spliced products without the effects of differential mRNA stability.

We first compared the splicing efficiencies (SE; the relative number of reads from spliced and unspliced RNAs) in the Prp8^WT^- and Prp8^Q1594A^-containing strains (**Fig. 6A**). Among all the detected introns (*N* = 331), 67% (202) exhibited significantly decreased SE in the presence of Prp8^Q1594A^, whereas only 5 genes showed increased SE (**Fig. 6A**). There was no apparent difference in the impact on ribosomal protein gene (RPGs) splicing versus non-RPGs, indicating that Prp8^Q1594A^ affects splicing generally and not necessarily related to transcript abundance or biological function.

**Figure 6.**
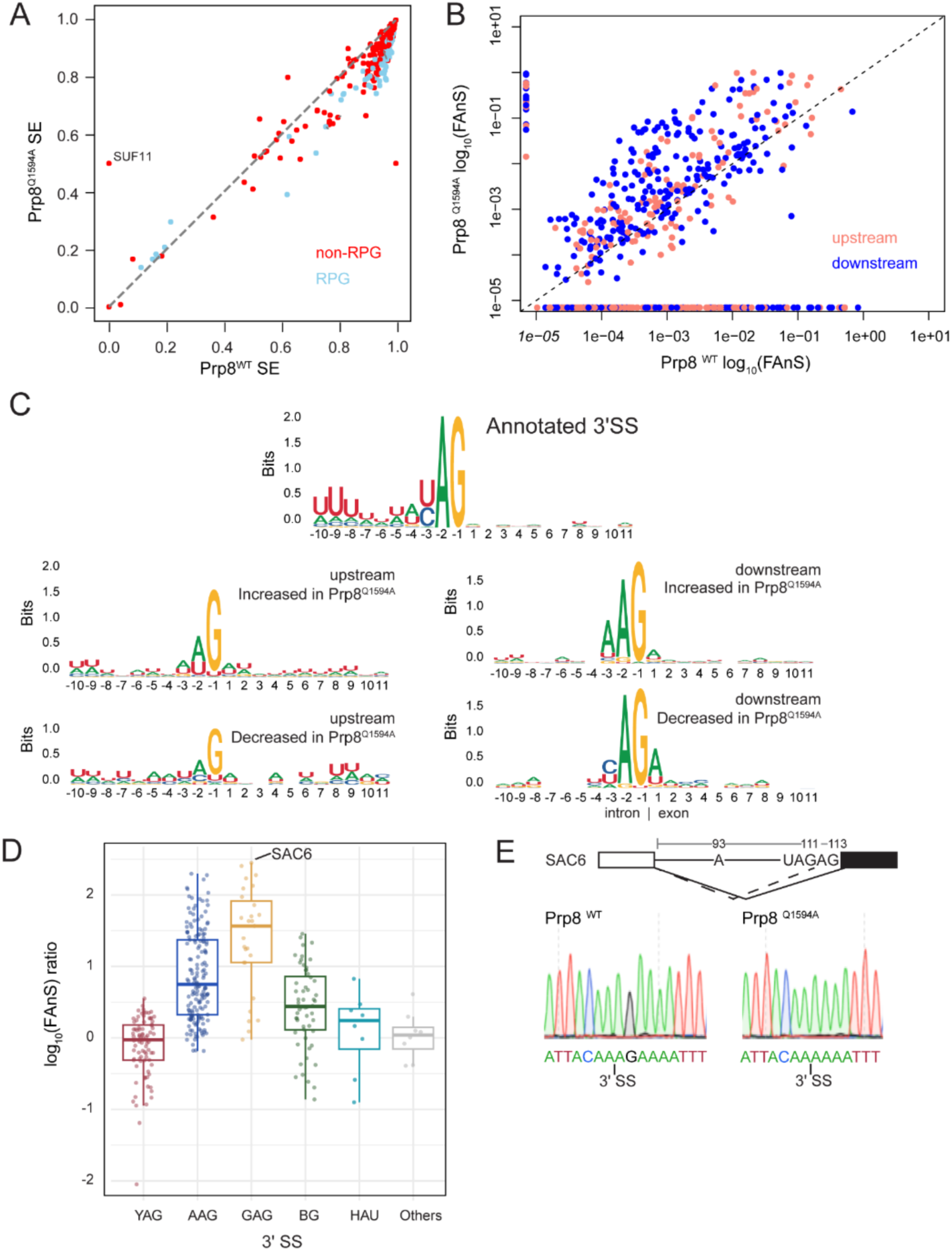
RNA-seq Analysis of Prp8^Q1594A^ Effects on Splicing. **(A)** Scatter plot comparing splicing efficiency (SE) between yeast strains with Prp8^WT^ or the Q1594A mutant. Points represent transcripts colored according to their identities as either ribosomal protein genes (RPG, blue) or non-ribosomal protein genes (non-RPG, red). Points below the diagonal represent lower SE in the mutant strain. SUF11 (tRNA-Pro) is noted as it shows increased splicing efficiency, but this intron is not processed by the spliceosome. **(B)** Differences in Fraction of annotated splicing (log_10_(FAnS)) between Prp8^WT^ and the Q1594A mutant. Alternative 3’SS Events are categorized based on the positions of the alternate site upstream (pink) or downstream (blue) relative to canonical site. Points above the diagonal indicate increased usage of the alternate site in the mutant strain. **(C)** Sequence logos of nucleotide preference around alternative 3’SS, categorized into the annotated site and upstream or downstream alternative sites. Sites increased in Prp8^Q1594A^ are those with log_10_(FAnS) ratios > 0 and those with ratios < 0 have decreased usage. **(D)** Boxplot comparison of alternative 3’SS usage frequencies expressed as ratios of the log_10_(FAnS) values for different sequences. Those with log_10_(FAnS) ratios > 0 have increased usage in the mutant strain **(E)** Representative sequencing results of cDNAs obtained from *SAC6* mRNAs isolated from yeast with either Prp8^WT^ or Prp8^Q1594A^.

To further explore how Prp8^Q1594A^ affects splice site selection, we then analyzed alternative splice junctions. We used rMATs to detect alternative splicing events and found that the most common type of alternative splicing event in both Prp8^WT^ and Prp8^Q1594A^ strains was use of an alternate 3’SS (**Sup. Fig. S5**). Of the 173 alternative 3’SS events identified in the Prp8^Q1594A^ strain, 147 (∼85%) were also identified in the Prp8^WT^ strain. We then analyzed the alternative 3’SS events by calculating Fraction of Annotated Splicing (FAnS) values for all alternative splice junctions. The FAnS value represents the relative abundance of an alternative splicing event compared to the canonical, annotated spliced isoform. Most alternative 3’SS events had larger FAnS values in the Prp8^Q1594A^-containing strain compared with Prp8^WT^ (**Fig. 6B**, points above dotted line). For the sites with increased usage, they were located both upstream and downstream of the annotated 3’SS with no apparent preference for alternative 3’SS location. In sum, these data indicate that while the Prp8^Q1594A^ mutation does not result in creation of a large number of new alternative 3’SS, it does significantly increase the use of alternative or cryptic 3’SS that are poorly utilized by Prp8^WT^.

To determine additional characteristics influencing alternative 3’SS selection, we calculated the log_10_ value for the ratio of the FAnS score for each site between the Prp8^Q1594A^ and Prp8^WT^ strains. We then analyzed sequence features of 3’SS with increased (log_10_FAnS ratio > 0) and decreased usage (log_10_FAnS ratio < 0) with the Prp8^Q1594A^ mutant (**Fig. 6C**). Annotated 3’SS are characterized by a strong YAG consensus motif. The alternate 3’SS which showed increased usage with Prp8^Q1594A^ could deviate from this at each position. While a guanine was still greatly preferred at the −1 position, Prp8^Q1594A^ enabled use of 3’SS with more variation at the −2 and −3 positions upstream of the annotated 3’SS. Sites increased in Prp8^Q1594A^ located downstream of the annotated 3’SS also showed more variation that then annotated sites but less so than those located upstream. These downstream sites tended to favor AAG sequences, which are spliced relatively well in yeast containing either Prp8^WT^ or the Q1594A mutant (**Fig. 2**). Alternate 3’SS less frequently used by the Prp8^Q1594A^ mutant showed strong variation if they were located upstream of the annotated site but were more similar to the YAG consensus if located downstream.

### Prp8^Q1594^ Changes Splice Site Competition in Endogenous RNAs

We next analyzed the alternate 3’SS based on their specific sequences (**Fig. 6D**, **Sup. Fig. S5B**). While the Prp8^Q1594A^ mutation did not change usage of 3’SS harboring canonical YAG sequences, it increased usage of sites with nonconsensus sequences. The greatest increases were observed for AAG and GAG 3’SS. For example, we noticed that the SAC6 gene encodes a UAGAG sequence at the 3’ end of its intron (**Fig. 6E**). The annotated 3’SS occurs after the consensus UAG sequence. We observed preferential use of this site in the Prp8^WT^ strain by RNA-Seq (**Sup. Fig. S5**) and confirmed its use by Sanger sequencing of the RT-PCR product (**Fig. 6E**). However, the GAG site was activated in the presence of Prp8^Q1594A^ and was detected in sequencing results obtained from cDNAs isolated from the Prp8^Q1594A^-containing strain (**Fig. 6E**, **Sup Fig. S5**). Average FAnS values of alternate 3’SS sequences indicated that the distribution of sequence preferences for alternate sites differs between Prp8^WT^ and Prp8^Q1594A^-containing strains, with notable changes in the values for GAG and AAG sites (**Sup. Fig. S5B**). While we observed significant increase in usage of a UCG 3’SS in the ACT1-CUP1 reporter (**Fig. 2**), we were unable to identify any UCG 3’SS with increased usage in Prp8^Q1594A^. We note that a limitation of our data is that very low numbers of reads mapped to potential UCG sites in general (often <5) and different mapping approaches may be able to facilitate identification of these or other unusual splicing events (Roy et al. 2023).

Finally, we identified several other transcripts with competing 3’SS and analyzed the associated RNA-Seq data. Yeast have two identified transcripts that contain naturally-occurring NAGNAG type 3’SS: *ECM33* (AAGCAG) and an intron located in the 5’ untranslated region (UTR) of the *BMH2* RNA (CAGAAG) (Schreiber et al. 2015). We examined our RNA-Seq data to determine if mutation of Prp8 would influence NAGNAG site usage in these RNAs. In the presence of Prp8^WT^, the consensus CAG site was used for both *ECM33* and *BMH2* (**Sup. Fig. S6**). However, approximately equal numbers of junction reads were present for the consensus and nonconsensus (AAG) sites for both transcripts with Prp8^Q1594A^. We also analyzed junction reads from the *APE2* and *DMC1* RNAs, which both contain BPS-proximal cryptic AAG 3’SS and canonical BPS-distal YAG 3’SS (**Sup. Fig. S7**) (Meyer et al. 2011). For both transcripts, we observed relatively few spliced-junction reads for all sites. Nonetheless, we noted that the Prp8^Q1594A^ mutation resulted in a switch in relative read count abundance. For Prp8^WT^, reads from the consensus site junctions outnumbered those from the cryptic sites. The opposite was true for reads from the Prp8^Q1594A^-containing strain. Together, these results indicate that the Prp8 α-finger can control mRNA isoform production by influencing how 3’SS compete with one another.

## DISCUSSION

Accurate selection of the 3’SS is essential for correct pre-mRNA splicing and gene expression fidelity (Will and Luhrmann 2011). While early studies established that Prp8 plays a central role in splicing catalysis and 3’SS positioning (Galej et al. 2016; and reviewed in Grainger and Beggs 2005), the observation of an interaction between the α-finger domain of Prp8 and the 3’SS in recent structures of post-catalytic (P complex) spliceosomes suggested that Prp8 influences exon ligation by direct interaction with the 3’SS to enforce SS identity (Bai et al. 2017; Liu et al. 2017; Wilkinson et al. 2017). In this study, we demonstrate that a conserved residue located with the Prp8 α-finger domain (Q1594) can control 3’SS usage *in vivo.* Replacement of the glutamine with alanine increases splicing efficiencies for non-YAG sites in both reporter pre-mRNAs and endogenous transcripts. Consequently, this can change how 3’SS compete with one another during splicing and alter mRNA isoform production. Genetic interactions between Prp8^Q1594A^ and Prp22 mutants suggest that perturbation of α-finger domain interactions with the 3’SS and/or the Prp22 CTD influences Prp22-dependent activities. This argues for a mechanism in which splice site recognition by Prp8 controls activity of and proofreading by Prp22.

### Context Dependence of 3’ SS Usage

How 3’SS compete with one another and what ultimately dictates use of the preferred site remain outstanding questions in understanding the molecular basis of alternative splicing regulation. In our experiments, we exploited the ability of the Prp8^Q1594A^ mutant to increase usage of non-YAG sites to study competing 3’SS in different contexts. Much of our data are consistent with selection occurring by a leaky scanning-like mechanism in the 5’→3’ direction (Semlow and Staley 2012). This is consistent with the 3’SS usages we observed in ACT1-CUP1 reporters harboring NAGNAG sites as well as in several endogenous transcripts including *SAC6*, *ECM33*, *BMH2*, *APE2*, and *DMC1*. In these cases, the consensus site predominates when yeast contain Prp8^WT^ but nonconsensus sites become more competitive with Prp8^Q1594A^. It is interesting to note that this includes the CAGAAG site in the *BMH2* RNA where the distal AAG site becomes competitive with the proximal CAG, consensus site when Prp8^Q1594A^ is present (**Sup. Fig. S6**). This can likely be explained by some degree of Prp22-dependent proofreading and 3’SS sampling (*i.e.,* binding, release, and re-binding) occurring even when a consensus site is present (Abelson et al. 2010; Semlow et al. 2016).

Some of our results, however, suggest additional complexities in 3’SS selection. First, we observed cases of cryptic 3’SS activation occurring *downstream* of the canonical site in the RNA-Seq data (**Fig. 6B**). These sites tended to have relatively strong, AAG consensus 3’SS sequences and competition may be driven in part by the strengths of these sites. We do not know if other factors such as RNA structures, sequences, or even use of alternate branch sites influence activation of these sites. Further studies are needed to disentangle these potential effects. Second, it is interesting to note that we did not observe any mRNA production by primer extension with ACT1-CUP1 reporters using UAY/AG sites in yeast with Prp8^Q1594A^. In these cases, the inclusion of pyrimidines activates overlapping YAGs site to create out-of-frame mRNAs when Prp8^WT^ is present (**Fig. 2**, **Sup. Fig. S2**). However, the use of these sites are prevented with Prp8^Q1594A^. One possibility is that the UAY 3’SS becomes “stuck” or the overlapping YAG cannot be sampled by Prp22-dependent proofreading when this Prp8 mutation is present. There is precedent for spliceosomes being able to accommodate UAY sequences in the active site: a cryo-EM structure was obtained of a human C* complex (a state post-5’SS cleavage and rearrangement but prior to exon ligation) with a likely “AC” sequence in the active site (Dybkov et al. 2023). In this case, the spliceosome was stalled at the C* state and did not (or could not) proceed to exon ligation. Finally, we also saw activation of cryptic AAG sites in the *APE2* transcript that are believed to be normally obscured, to some degree, by a stem loop structure (Meyer et al. 2011). Since we grew and collected Prp8^WT^ and Prp8^Q1594A^ under identical temperature conditions, our results suggest that the thermal stability of this intron structure alone cannot account for *in vivo* splice site usage. It is likely the combination of this structure and the molecular properties of the assembled spliceosome that dictate to what degree these cryptic sites may be used.

### Evidence for Conformational Change of the Prp8 α-finger Domain

Our data strongly support a critical role for the Prp8 α-finger in 3’SS selection. To study the function of this domain further, we analyzed its conformation in models of C* (pre-exon ligation), P (post-exon ligation), and ILS (intron lariat spliceosome; post mRNA release) yeast spliceosomes obtained by cryo-EM. We noticed that the α-finger was only modeled in P complex structures that also contained a bound 3’SS (**Fig. 7A**). Together with Max Wilkinson, we recently published a high-resolution structure of the yeast spliceosome P complex (Senn et al. 2024). In that data set, we noted that ∼29% of the particles represented a unique state of P complex (called State III, **Fig. 7B**, **C**). In the state III complex, the 3’SS was absent from the active site, several 2^nd^ step factors could not be observed (Fyv6, Prp18, and Slu7), and cryo-EM density was lacking for much of the α-finger.

**Figure 7.**
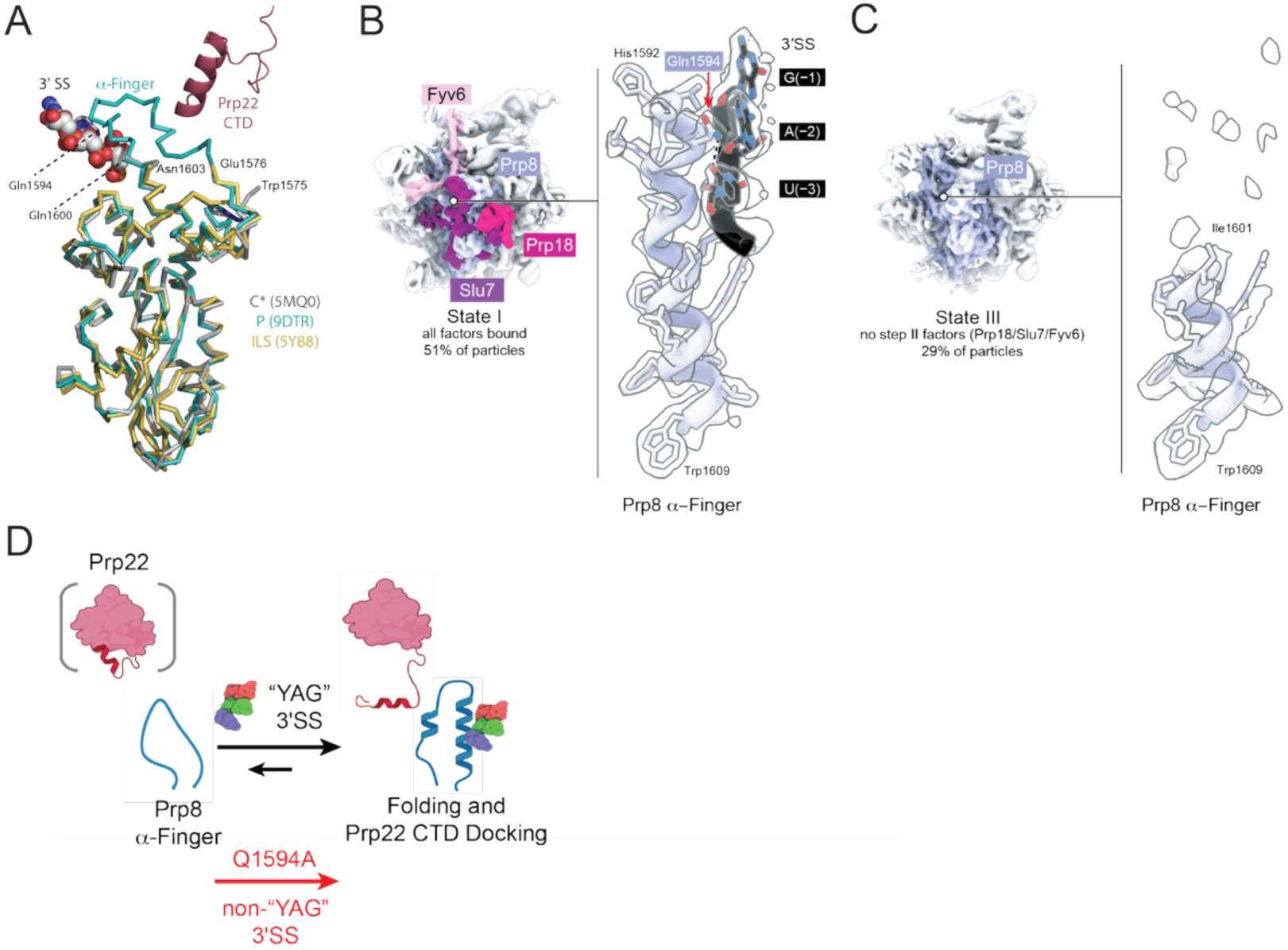
Model for Regulation of Prp22 Activity by Prp8 α-Finger Domain Dynamics. **(A)** Overlay of the Prp8 α-finger region from cryo-EM structures of C* (grey, 5MQ0), P (blue, 9DTR), and ILS (yellow, 5Y88) complex spliceosomes. The α-finger could only be modeled in the P complex structure, in which the 3’SS and Prp22 CTD were also present. Figure generated using PyMOL (Schrödinger, LLC). **(B, C)** Comparison of cryo-EM densities for State I and State III P complex spliceosomes from (Senn et al. 2024). Clear density for the α-finger was not observable in the State III complexes, which also lacked the 2^nd^-step factors Fyv6, Prp18, and Slu7. State III complexes also lacked observable densities for the 3’SS and Prp22 CTD. Figure generated using ChimeraX (Meng et al. 2023). **(D)** In this model, binding of the Prp22 CTD is mutually exclusive with an unfolded conformation of the Prp8 α-finger. Binding of a potential 3’SS facilitates folding of the α- finger and docking of the Prp22 CTD near the active site (or *vice versa*). The Prp8^Q1594A^ mutation promotes folding of the α-finger and Prp22 CTD docking when non-YAG 3’SS are present. Figure created with BioRender software.

Together, these observations suggest that the α-finger is conformationally dynamic and may only adopt a helical conformation when a 3’SS has docked into the active site and 2^nd^-step factors are present. We do not know if folding of the α-finger permits 3’SS docking, if presence of a competent 3’SS causes folding, or if docking and folding are concerted events. However, we favor a model in which the α-finger is folded prior to exon ligation. This is based on the cryo-EM structure of the human C* complex containing an “AC” dinucleotide docked into the active site. In this structure, the α-finger is folded and the 5’ exon has not yet been ligated at the 3’SS (Dybkov et al. 2023). This suggests that α-finger folding precedes bond formation, whereas our state III P complex structure indicates that unfolding of the α-finger precedes ILS structure formation and mRNA release. In summary, we propose that folding of the α-finger is a necessary step in splicing that occurs prior to exon ligation and that unfolding occurs to facilitate mRNA release and P complex remodeling to the ILS conformation.

### A Model for Regulation of Prp22 Activity by the Prp8 α-Finger

In addition to whether the Prp8 α-finger could be modeled, we also noted that the ability to model the Prp22 CTD as inserted into the spliceosome active site also correlated with presence of the 3’SS and folding of the α-finger. The Prp22 CTD was observed in the state I (3’SS docked) but not that state III yeast P complex structure (3’SS undocked) (Senn et al. 2024), it was modeled in the human C* structure containing the “AC” 3’SS mimic (Dybkov et al. 2023), and it was also modeled in the human P complex structure containing a bound 3’SS and partially ordered α-finger (Fica et al. 2019). Since the ultimate selection of the 3’SS likely depends on coupling of the exon ligation step to Prp22-dependent spliceosome remodeling and mRNA release (Mayas et al. 2006; Semlow et al. 2016), it is possible that folding of the α-finger is coupled to Prp22 CTD insertion into the active site. We propose that when the α-finger is unfolded, it sterically blocks the Prp22 CTD from inserting into the core of the spliceosome (**Fig. 7D**). When it is folded and a potential 3’SS is bound to the active site, the Prp22 CTD can be accommodated.

This model would directly correlate Prp22 binding with conformation of the spliceosome active site. It would also provide a means for regulating Prp22-dependent activities. By preventing Prp22 CTD domain insertion, it is possible that Prp22 cannot interact with the mRNA or the spliceosome in a manner to facilitate mRNA release. Thus, folding of the α-finger just prior to exon ligation may signal that the spliceosome is ready to engage with Prp22. In our model, the Prp8^Q1594A^ mutation facilitates folding of the α-finger on non-consensus, non-YAG substrates to permit Prp22 CTD docking and exon ligation (**Fig. 7D**).

An important limitation of this model is that we do not know yet how docking of the Prp22 CTD influences Prp22 ATPase or translocation activities themselves. Does CTD docking stimulate, inhibit, or not change ATP hydrolysis? We favor a model in which Prp22 CTD docking inhibits ATP hydrolysis since unfolding of the α-finger and CTD dissociation could provide a mechanism for signaling that the spliceosome is ready to release a spliced mRNA product. This model could also provide a mechanism to explain proofreading by Prp22: slow (or no) folding of the α-finger and failure to insert the CTD due to non-splicing competent 3’SS may fail to “switch off” the Prp22 ATPase, result in active site disruption prior to transesterification, and permit sampling of alternate 3’SS. In this case, a potential mechanism for suppression of the *ts* phenotype due to Prp8^Q1594A^/Prp22^G810A^ mutations by Prp22^I1133R^ (**Fig. 5**) could be due to weaker binding of the Prp22 CTD and upregulated Prp22 ATPase activity despite the Prp22^G810A^ mutation. This may counter losses in splicing fidelity due to the combined effects of the Prp8^Q1594A^ and Prp22^G810A^ mutations. Future work may be able to test these hypotheses by studying biochemical activities of Prp22 ± CTD, in-depth transcriptomic analysis of Prp8 and Prp22 mutant-contianing yeast strains, and/or by capturing spliceosome C* complexes formed on non-YAG sites or P complexes during mRNA release by cryo-EM.

## MATERIALS AND METHODS

### Yeast strain and plasmid construction

Yeast strain construction, molecular cloning, and site-directed mutagenesis were performed as previously described (Lipinski et al. 2023). The PRP8 shuffle strain yAAH0117 (yJU75) lacking genomic PRP8 was generously provided by Christine Guthrie and provided on a URA3-based plasmid (Umen and Guthrie 1996). The Prp8 mutant plasmids, Prp22 mutant plasmids, and ACT1-CUP1 reporters were prepared by PCR-based site-directed mutagenesis. The UPF1 and FYV6 genes were deleted by replacement with a kanamycin resistance cassette (KanMX) and a nourseothricin resistance cassette (NatMX4), respectively, through homologous recombination(Goldstein and McCusker 1999). Gene deletion was confirmed by colony PCR.

### RNA isolation

Yeast cultures were grown in YPD medium to an OD600 of 0.5–0.8. Cells equivalent to 15 OD600 units were collected by centrifugation (4000 rpm, 5 min, 4°C), resuspended in ice-cold RNase-free water or PBS, and centrifuged again to remove the supernatant. Cell pellets were resuspended in 1.5 mL ice-cold TRIzol and disrupted using a Disruptor Genie for 1 min, followed by incubation on ice (1 min). This cycle was repeated three times to ensure complete lysis. The lysate was then transferred to a Phasemaker™ tube (Invitrogen) and incubated at room temperature for 3 min. Subsequently, chloroform (300 µL) was added, and the mixture was vigorously shaken for 30 s, incubated at room temperature for 5 min, and centrifuged at 2500 × g for 5 min at 4°C. The resulting aqueous phase was carefully collected into a pre-chilled microcentrifuge tube for downstream RNA purification.

RNA purification was carried out using the Monarch® RNA Cleanup Kit (NEB) according to the manufacturer’s instructions. Briefly, an equal volume of 100% ethanol was added to the TRIzol-isolated aqueous phase and mixed gently. The mixture was applied to a Monarch RNA Cleanup column in several steps, centrifuged at 10,000 × g for 1 min at 4°C, and the flowthrough was discarded after each step. The column was washed twice, followed by a final centrifugation to remove residual ethanol. RNA was eluted in 30 µL of RNase-free water. To remove genomic DNA contamination, TURBO DNase (Thermo Fisher Scientific) treatment was performed. DNase buffer and enzyme were added to the RNA sample and incubated at 37°C for 20 min. TURBO DNase inactivation reagent was then added, gently mixed by flicking, and incubated at room temperature for 5 min. The mixture was centrifuged at 10,000 × g for 1.5 min at 4°C, and the supernatant containing DNase-treated RNA was transferred to a fresh RNase-free tube.

RNA quality was assessed using a NanoDrop to determine concentration and purity. For RNA used in RNA-seq, RNA integrity and accurate quantification were further assessed using a Qubit fluorometer (Thermo Fisher Scientific) and TapeStation system (Agilent Technologies). Only high-quality RNA samples with minimal rRNA degradation, RNA integrity number (RIN) > 7.0, and A260/A280 ratios near 2.0 were selected for library preparation. RNA samples were stored at – 80°C until further use.

### Library preparation and RNA-seq

Library preparation was performed at the Gene Expression Center of the University of Wisconsin–Madison Biotechnology Center. mRNA was enriched from total RNA using poly(A) selection with the TruSeq Stranded mRNA Library Prep Kit (Illumina). Libraries were sequenced on an Illumina NovaSeq 6000 platform, generating paired-end reads.

### Bioinformatic analysis of RNA-seq database

RNA-Seq data analysis began with quality control of raw and trimmed reads using FASTQC (Andrews 2010). Adapter trimming and quality filtering were performed with fastp. Clean reads were mapped to the *Saccharomyces cerevisiae* reference genome (SacCer3, Ensembl R64-1-1) using STAR (Dobin et al. 2013). After the first round of alignment, novel splice junctions were extracted and filtered to remove potential false positives. The filtered novel splice junctions were then used as additional input in a second round of STAR alignment to improve junction detection (Senn et al. 2024). Following alignment, bam files were indexed using SAMtools (Li et al. 2009) and read counts for splice junctions within genes were quantified using featureCounts (Liao et al. 2014). Splicing efficiency (SE) for each intron was calculated as previously described (Roy et al. 2023).

Canonical branch points (BP) were annotated based on the Ares Intron Database (Grate and Ares 2002). Canonical 5’ and 3’ splice sites (SS) were annotated based on the junctions with the highest read counts. All junctions were filtered to retain only those present in all six RNA-seq samples (triplicates for Prp8^WT^ and Prp8^Q1594A^ strains). Alternative splice junctions were categorized based on splicing patterns, including those that shared a canonical 5’ SS but had an alternative 3’ SS, or those that shared a canonical 3’ SS but had an alternative 5’ SS. FAnS was calculated by dividing the number of junction reads for an alternative 3’ SS that shares a canonical 5’ SS by the total number of junction reads for canonical 5’ and 3’ SS as describe before (Roy et al. 2023). Alternative splicing events were classified using rMATS (Shen et al. 2014; Wang et al. 2024) and MASER, and visualized using the Integrative Genomics Viewer (IGV). For rMATS, the flag “--novelSS” was used to detect novel splice sites.

### Primer extension

Primer extension was performed to detect RNA extension products from the ACT1–CUP1 reporter gene and U6 snRNA, as previously described (Carrocci et al. 2017). Total RNA was extracted as described above. IR700 dye-labeled oligonucleotides (Integrated DNA Technologies) targeting ACT1–CUP1 (yAC6: /5IRD700/GGCACTCATGACCTTC) and U6 snRNA (yU6: /5IRD700/GAACTGCTGATCATGTCTG) were used. The primer mix contained 2 µM yAC6 and 0.4 µM yU6, along with dNTPs and 10 µg of total RNA. The mixture was heated at 65°C for 5 min and placed on ice for 5 min. Reverse transcription was performed with SuperScript III (Thermo Fisher Scientific) in the presence of RNase inhibitor, DTT, and the provided buffer, followed by incubation at 55°C for 1 h. Reactions were terminated by heating at 95°C, cooled on ice, and stored at –20°C until analysis.

Extension products were resolved on a 7% (w/v) denaturing polyacrylamide gel (8 M urea, 1× TBE). Gels were pre-run for 20 min, and samples were electrophoresed at 35 W for 1 h 20 min. Fluorescent signals were visualized using an IR Typhoon scanner (Cytiva).

### RT-PCR and cloning of SAC6 cDNA

RT-PCR was carried out using the Access RT-PCR System (Promega). 100 ng of total RNA was used per 25 µL reaction. *SAC6* transcripts were amplified using the primers SAC6-F (5′- ATGAATATTGTCAAATTACAA-3′) and SAC6-R (5′-GTCAATGGCTCTGAAC-3′). PCR products were visualized by ethidium bromide staining after gel electrophoresis, purified, and then cloned into plasmids using a TOPO^TM^ kit (Thermo Fisher Scientific). Plasmids were transformed into Top10 *Escherichia coli* cells, plasmids isolated by miniprep, and Sanger sequenced;.

### Yeast temperature growth assay

Yeast cultures were grown overnight in YPD medium to stationary phase, then adjusted to OD600 = 0.5 in 10% (v/v) glycerol. Three consecutive 1:10 serial dilutions were performed by transferring 100 µL of culture into 900 µL of diluent, resulting in final dilutions of 1:10, 1:100, and 1:1000. Diluted cultures were then spotted onto YPD agar plates and incubated at 16°C, 23°C, 30°C, or 37°C for the time specified in the corresponding figures before imaging.

### ACT1-CUP1 assay

To assess splicing-dependent copper tolerance, yeast strains harboring ACT1–CUP1 reporter plasmids were cultured in selective dropout media. Strains carrying the Prp18^V191A^ reporter were grown in -Leu/-Ura medium, while others were cultured in -Leu medium to maintain plasmid selection. Diluted cultures were spotted onto selective dropout plates supplemented with 0-2.5 mM CuSO_4_ (Lesser and Guthrie 1993; Carrocci et al. 2018) and incubated at 30°C before imaging.

## DATA AVAILABILITY

Data are available via the NCBI Sequence Read Archive (BioProject ID PRJNA1236378).

## SUPPORTING INFORMATION

This manuscript contains supporting information, including supplementary figures and tables. These materials can be accessed online alongside the full text of the article.

## AUTHOR CONTRIBUTIONS

YL, AAH: project conceptualization; YL, JCP, AAH: media and reagent preparation; YL: data acquisition and analysis; YL, AAH: writing and editing of the manuscript.

## ACKNOWLEDGEMENTS

We thank Drs. Karli Lipinski and Kathy Senn for help with RNA-Seq data analysis and ACT1-CUP1 assays. We thank Dr. Beate Schwer for Prp18 plasmids. We thank Dr. Max Wilkinson for help in making Figure 7 as well as insightful discussions. We thank Drs. Jon Staley and Guillaume Chanfreau for helpful discussions and feedback on the manuscript.

