## Supplemental Figures and Tables for "Control of 3’ Splice Site Selection in *S. cerevisiae* by a Highly Conserved Amino Acid within the Prp8 α-finger Domain"

**CORRESPONDING AUTHOR:**

**CONTENTS:**

Supplemental Figures S1-S7

Supplemental Tables S1 and S2

Supplemental References

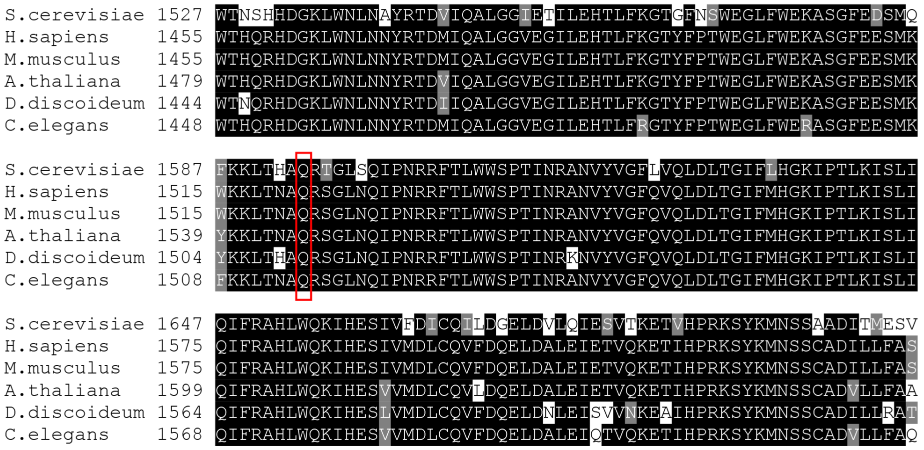


**Supplementary Figure S1. Multiple sequence alignment of the Prp8 $\alpha$-finger region from different species.** The red box marks yeast Q1594. Sequences were aligned and visualized using Clustal Omega (Madeira et al. 2024) and Boxshade (Albà 2000).

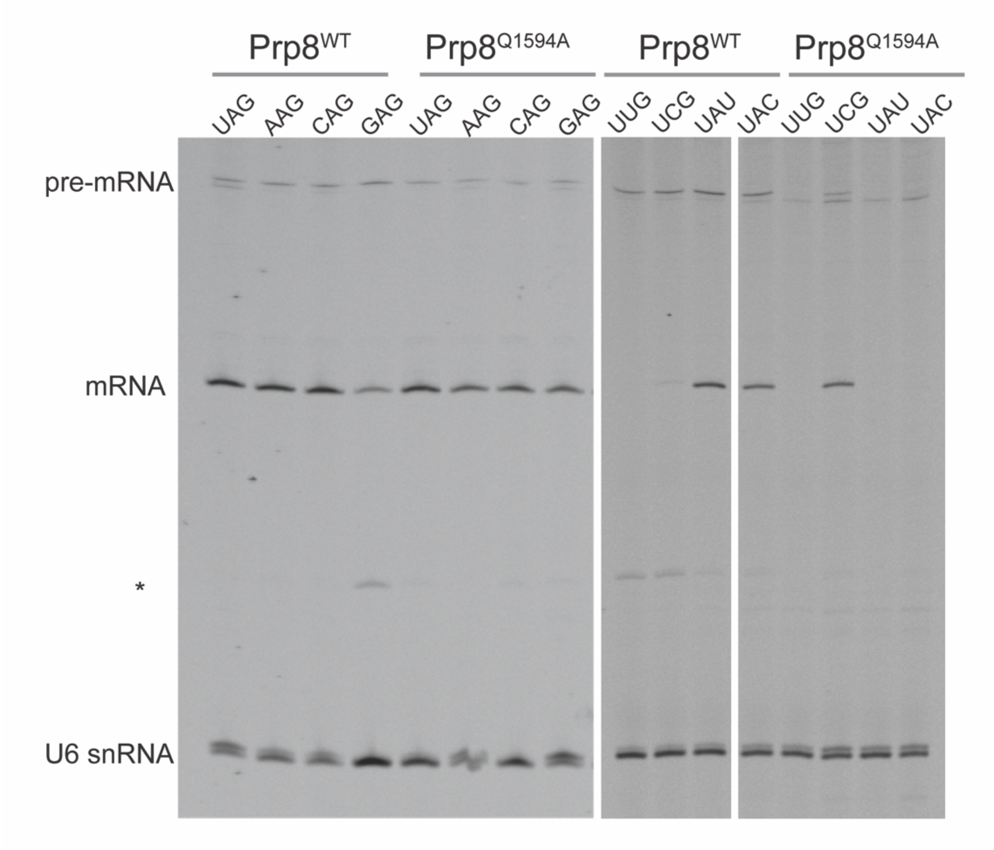


**Supplementary Figure S2. Primer extension analysis of ACT1-CUP1 reporter RNAs in the presence of Prp8$^{WT}$ or Prp8$^{Q1594A}$**. White bars indicate that the experiments were run on different gels or intervening lanes removed to generate this figure. Note that bands corresponding to mRNAs are observed for Prp8$^{WT}$ with the UAU and UAC reporters despite the low copper tolerances observed (see **Fig. 2**). This is likely due to splicing at a flanking "AG" site (UA<u>U/AG</u>; UA<u>C/AG</u>) created by the substitution at the normal 3'SS and a shift in the protein reading frame. Also note that these mRNAs are not observed if the same reporters are present with Prp8$^{Q1594A}$.

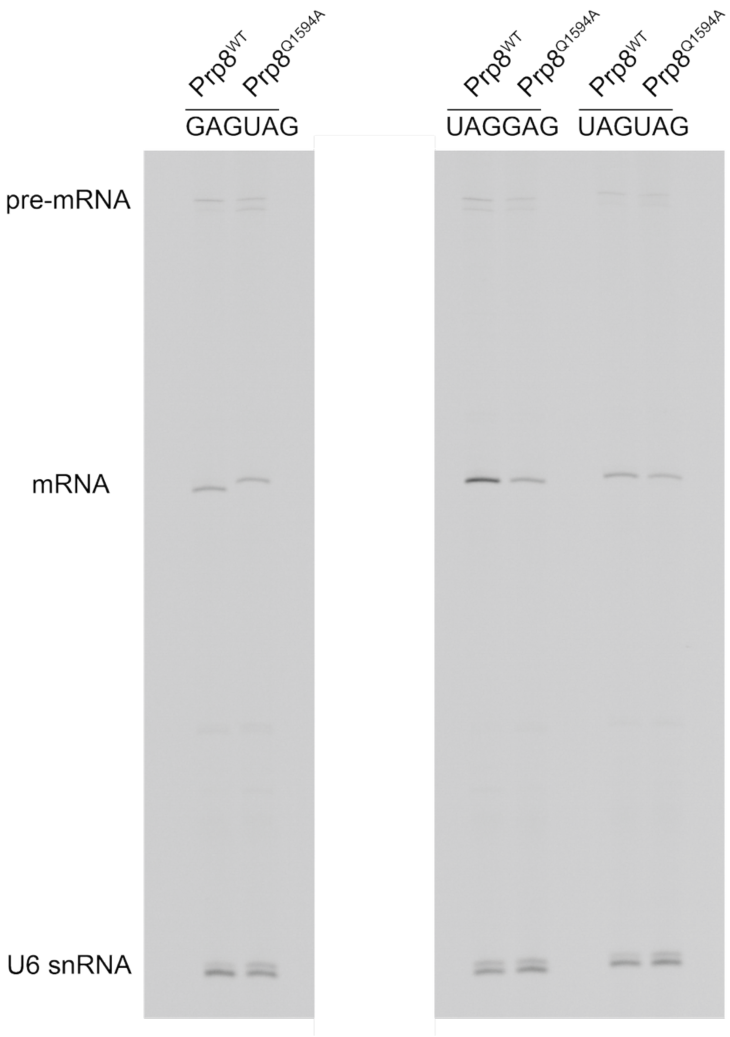


**Supplementary Figure S3. Primer extension analysis of ACT1-CUP1 reporters with competing 3'SS.** This is the full image corresponding to the cropped region shown in **Fig. 3B**. Intervening lanes have not been included and are covered by a white rectangle.

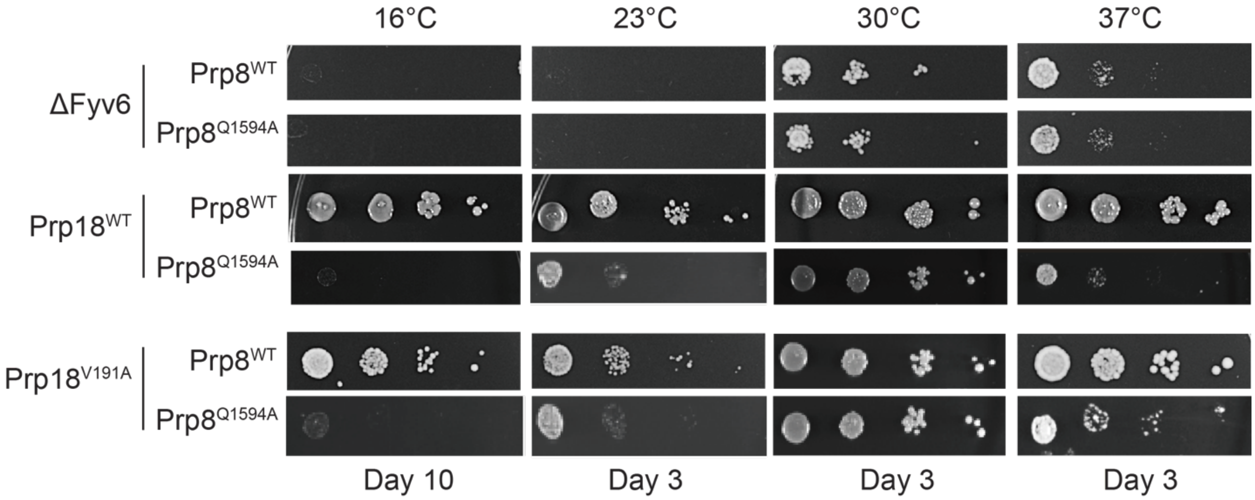


**Supplementary Figure S4. Yeast growth assays in Fyv6 deletion or Prp18 mutation backgrounds.** Growth assays comparing WT Prp8 and the Q1594A mutant at different temperatures in the absence of Fyv6 (∆Fyv6, YPD plates) or in the presence of WT or a dominant negative mutant of Prp18 (Prp18$^{V191A}$;-URA plates). Plates were incubated at the given temperatures and images collected on the noted days.

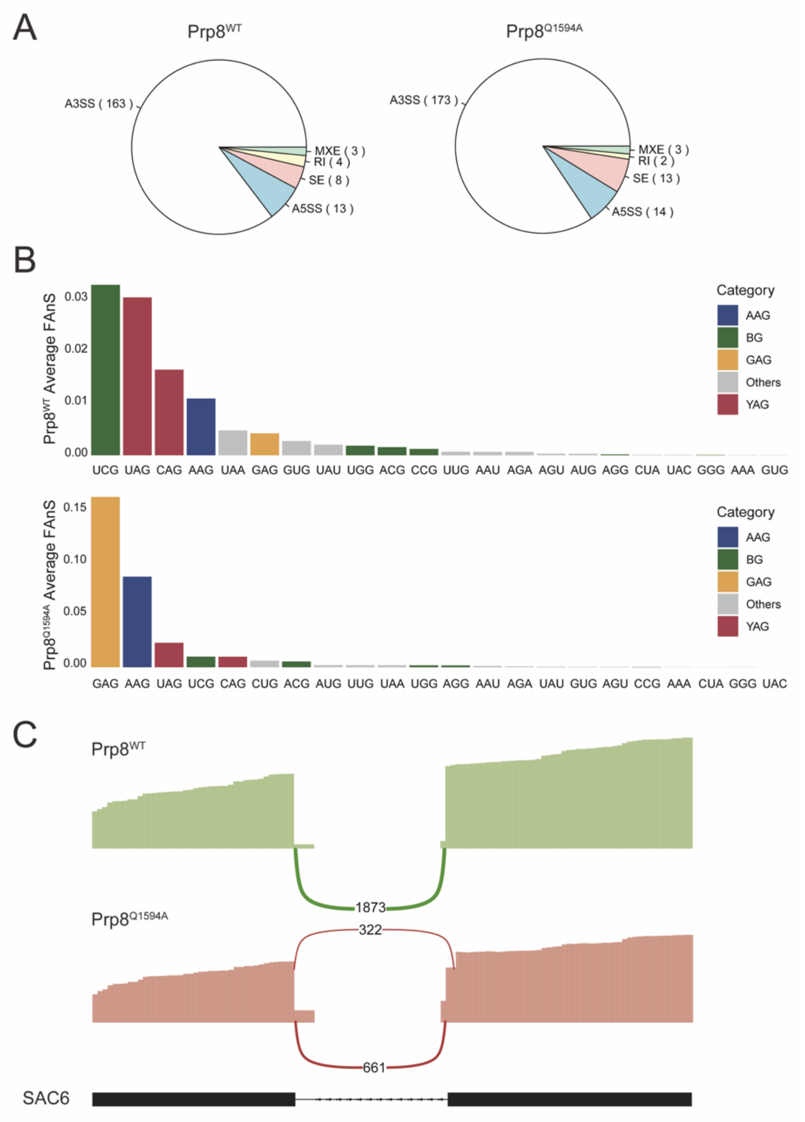


**Supplementary Figure S5. RNA seq analysis of Prp8$^{WT}$ or Prp8$^{Q1594A}$ yeast strains.** **(A)** Pie charts showing the numbers and types of alternative splicing events detected in the Prp8$^{WT}$ and Prp8$^{Q1594A}$ strains. **(B)** Bar graphs showing the average FAnS values for each 3'SS trinucleotide motif in Prp8$^{WT}$ (top) and Prp8$^{Q1594A}$ (bottom) for alternate 3'SS. Note that the Y axis is from 0.00-0.03 for Prp8$^{WT}$ but from 0.00-0.15 for Prp8$^{Q1594A}$. **(C)** Sashimi plot showing alternative 3'SS usage in the SAC6 gene. In the Prp8$^{WT}$ strain (green), the canonical distal 3'SS is predominantly used. In contrast, the Prp8$^{Q1594A}$ strain (red) shows increased usage of a proximal, cryptic 3'SS (~2:1 ratio of the cryptic to canonical site based on read counts).

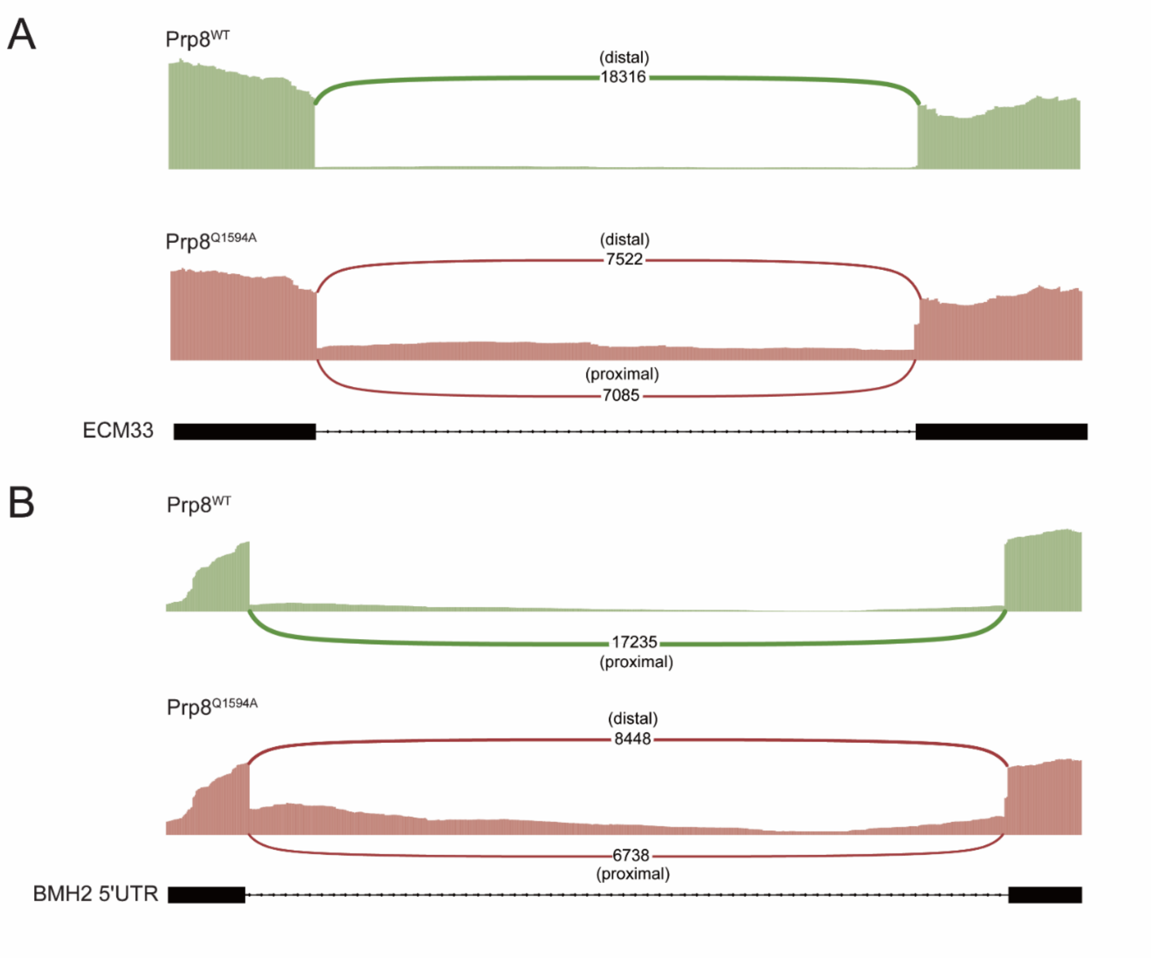


**Supplementary Figure S6. Sashimi plots showing alternative 3'SS usage in the ECM33 and BMH2 genes.** In the Prp8$^{WT}$ strain (green), the canonical 3'SS are predominantly used (distal in panel A and proximal in panel B). In contrast, the Prp8$^{Q1594A}$ strain (red) shows increased usage of cryptic 3'SS located proximal (panel A) and distal (panel B) to the BS.

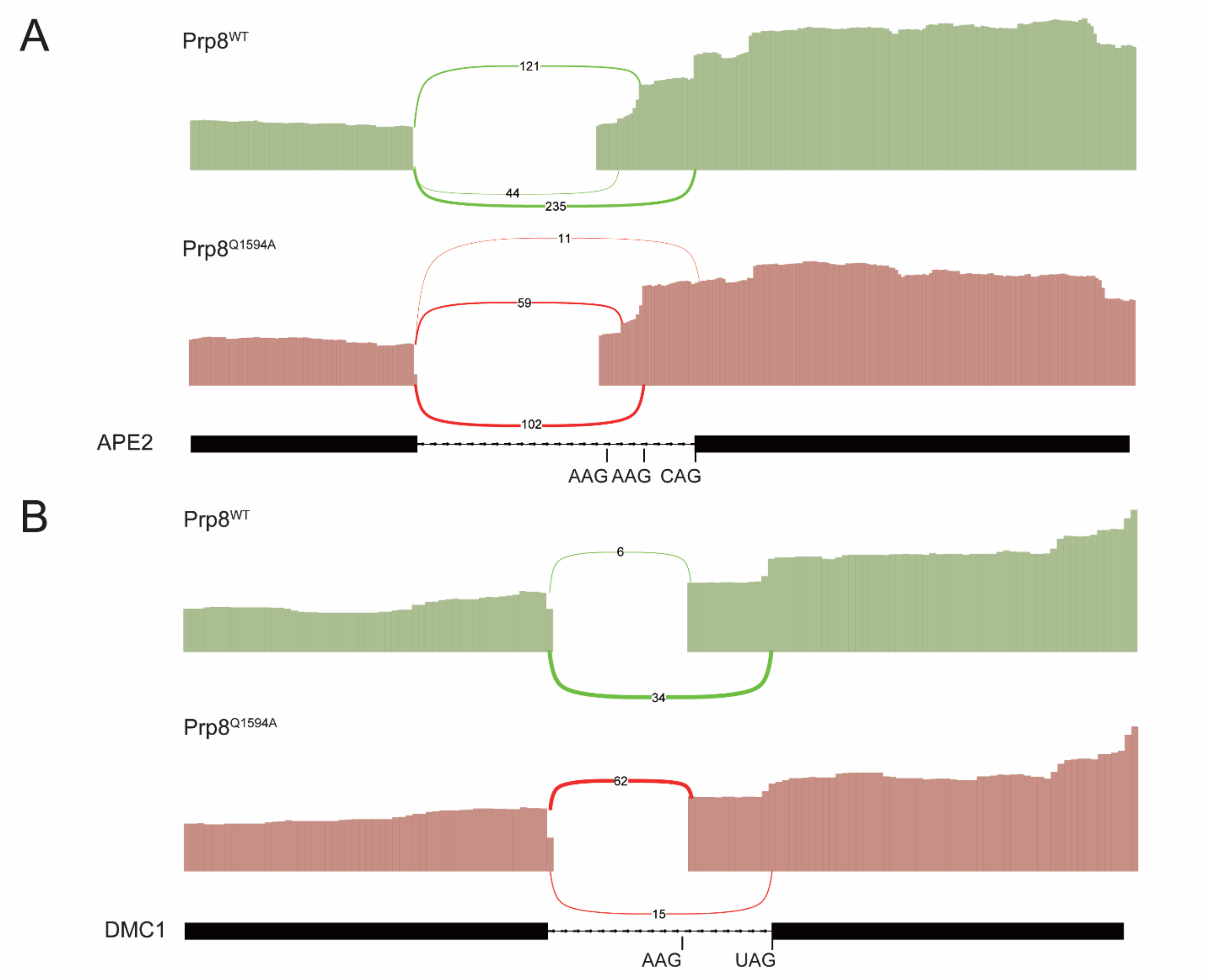


**Supplementary Figure S7. Sashimi plots showing alternative 3'SS usage in the APE2 and DMC1 genes. (A)** When Prp8$^{WT}$ is present, the canonical CAG site is most frequently used in the *APE2* RNA relative to two other AAG sites at a ratio of read counts of ~ 1:2.8:5.3 (going from most proximal to the BS to the most distal). When Prp8$^{Q1594A}$ is present, this ratio changes to 1:1.7:0.19. (**B**) For the *DMC1* RNA, we observed more read counts for usage of the canonical UAG site relative to the cryptic AAG site (ratio of 1:5.7) with WT Prp8. This ratio is nearly flipped when the Prp8$^{Q1594A}$ mutant is present (ratio of 1:0.25). Note that for both *APE2* and *DMC1* RNAs, we observed relatively few read counts that spanned exon-exon junctions.

**Supplementary Table S1.** Yeast strains used in this study.

| Strain | Genotype | Description | Reference |
|---|---|---|---|
| yAAH0117 | ade2, cup1delta:ra3 his3 leu2 lys2 prp8delta:lys2 trp1 pJU169:PRP8(URA) | Prp8WT strain | Umen and Guthrie, Genetics, 1996 |
| yAAH3853 | yAAH0117 + pAAH1440 | pJU169:PRP8_WT(TRP) | This work |
| yAAH3527 | yAAH0117 + pAAH1627 | pJU169:PRP8_Q1594A(TRP) | This work |
| yAAH3817 | ade2, cup1delta:ra3 his3 leu2 lys2 prp8delta:lys2 trp1 pJU169:PRP8(URA)UPF1::NATR | PRP8(URA) UPF1 deletion | This work |
| yAAH3605 | yAAH3817 + pAAH1440 | PRP8_WT(TRP) UPF1 deletion | This work |
| yAAH3818 | yAAH3817 + pAAH1627 | PRP8_Q1594A(TRP) UPF1 deletion | This work |
| yAAH3532 | yAAH3853 + pAAH0470 | PRP8_WT(TRP)ACT1-CUP1 WT | This work |
| yAAH3533 | yAAH3853 + pAAH0526 | PRP8_WT(TRP)ACT1-CUP1 U301G(3'ssGAG) | This work |
| yAAH3534 | yAAH3853 + pAAH1644 | PRP8_WT(TRP)ACT1-CUP1 U301A(3'ssAAG) | This work |
| yAAH3535 | yAAH3853 + pAAH1645 | PRP8_WT(TRP)ACT1-CUP1 U301C(3'ssCAG) | This work |
| yAAH3814 | yAAH3853 + pAAH1684 | PRP8_WT(TRP)ACT1-CUP1 3'ssGUGnew | This work |
| yAAH3544 | yAAH3527 + pAAH0470 | PRP8_Q1594A(TRP)ACT1-CUP1 WT | This work |
| yAAH3545 | yAAH3527 + pAAH0526 | PRP8_Q1594A(TRP)ACT1-CUP1 U301G(3'ssGAG) | This work |
| yAAH3546 | yAAH3527 + pAAH1644 | PRP8_Q1594A(TRP)ACT1-CUP1 U301A(3'ssAAG) | This work |
| yAAH3547 | yAAH3527 + pAAH1645 | PRP8_Q1594A(TRP)ACT1-CUP1 U301C(3'ssCAG) | This work |
| yAAH3816 | yAAH3527 + pAAH1684 | PRP8_Q1594A(TRP)ACT1-CUP1 3'ssGUGnew | This work |
| yAAH3541 | yAAH3853 + pAAH1752 | PRP8_WT(TRP)ACT1-CUP1 A302U(3'ssUUG) | This work |
| yAAH3542 | yAAH3853 + pAAH1753 | PRP8_WT(TRP)ACT1-CUP1 A302C(3'ssUCG) | This work |
| yAAH3543 | yAAH3853 + pAAH1754 | PRP8_WT(TRP)ACT1-CUP1 G303U(3'ssUAU) | This work |
| yAAH3548 | yAAH3853 + pAAH1755 | PRP8_WT(TRP)ACT1-CUP1 G303C(3'ssUAC) | This work |
| yAAH3549 | yAAH3527 + pAAH1752 | PRP8_Q1594A(TRP)ACT1-CUP1 A302U(3'ssUUG) | This work |

| | | | |
|---|---|---|---|
| yAAH3550 | yAAH3527 + pAAH1753 | PRP8_Q1594A(TRP)ACT1-CUP1 A302C(3'ssUCG) | This work |
| yAAH3551 | yAAH3527 + pAAH1754 | PRP8_Q1594A(TRP)ACT1-CUP1 G303U(3'ssUAU) | This work |
| yAAH3552 | yAAH3527 + pAAH1755 | PRP8_Q1594A(TRP)ACT1-CUP1 G303C(3'ssUAC) | This work |
| yAAH3530 | yAAH3853 + pAAH1729 | Prp8_WT(TRP) ACT1-CUP1 3'ss GAGUAG | This work |
| yAAH3529 | yAAH3527 + pAAH1729 | Prp8_Q1594A(TRP) ACT1-CUP1 3'ss GAGUAG | This work |
| yAAH3913 | yAAH3853 + pAAH1761 | Prp8_WT(TRP) ACT1-CUP1 3'ss UAGUAG | This work |
| yAAH3916 | yAAH3527 + pAAH1761 | Prp8_Q1594A(TRP) ACT1-CUP1 3'ss UAGUAG | This work |
| yAAH3914 | yAAH3853 + pAAH1762 | Prp8_WT(TRP) ACT1-CUP1 3'ss UAGGAG | This work |
| yAAH3917 | yAAH3527 + pAAH1762 | Prp8_Q1594A(TRP) ACT1-CUP1 3'ss UAGGAG | This work |
| yAAH3972 | ade2, cup1delta:ra3 his3 leu2 lys2 prp8delta:lys2 trp1 pJU169:PRP8(TRP) Fyv6::KAN | PRP8_WT(TRP) FYV6 deletion | This work |
| yAAH3973 | ade2, cup1delta:ra3 his3 leu2 lys2 prp8delta:lys2 trp1 pJU169:PRP8_Q1594A(TRP) Fyv6::KAN | PRP8_Q1594A(TRP) Fyv6 deletion | This work |
| yAAH3956 | yAAH3972 + pAAH0470 | PRP8_WT(TRP) FYV6_del ACT1-CUP1 WT | This work |
| yAAH3957 | yAAH3972 + pAAH0526 | PRP8_WT(TRP) FYV6_del ACT1-CUP1 U301G(3'ssGAG) | This work |
| yAAH3958 | yAAH3972 + pAAH1644 | PRP8_WT(TRP) FYV6_del ACT1-CUP1 U301A(3'ssAAG) | This work |
| yAAH3959 | yAAH3972 + pAAH1645 | PRP8_WT(TRP) FYV6_del ACT1-CUP1 U301C(3'ssCAG) | This work |
| yAAH3960 | yAAH3972 + pAAH1752 | PRP8_WT(TRP) FYV6_del ACT1-CUP1 A302U(3'ssUUG) | This work |
| yAAH3961 | yAAH3972 + pAAH1753 | PRP8_WT(TRP) FYV6_del ACT1-CUP1 A302C(3'ssUCG) | This work |
| yAAH3962 | yAAH3972 + pAAH1754 | PRP8_WT(TRP) FYV6_del ACT1-CUP1 G303U(3'ssUAU) | This work |

| | | | |
|---|---|---|---|
| yAAH3963 | yAAH3972 + pAAH1755 | PRP8_WT(TRP) FYV6_del ACT1-CUP1 G303C(3'ssUAC) | This work |
| yAAH3964 | yAAH3973 + pAAH0470 | PRP8_Q1594A(TRP) FYV6_del ACT1-CUP1 WT | This work |
| yAAH3965 | yAAH3973 + pAAH0526 | PRP8_Q1594A(TRP) FYV6_del ACT1-CUP1 U301G(3'ssGAG) | This work |
| yAAH3966 | yAAH3973 + pAAH1644 | PRP8_Q1594A(TRP) FYV6_del ACT1-CUP1 U301A(3'ssAAG) | This work |
| yAAH3967 | yAAH3973 + pAAH1645 | PRP8_Q1594A(TRP) FYV6_del ACT1-CUP1 U301C(3'ssCAG) | This work |
| yAAH3968 | yAAH3973 + pAAH1752 | PRP8_Q1594A(TRP) FYV6_del ACT1-CUP1 A302U(3'ssUUG) | This work |
| yAAH3969 | yAAH3973 + pAAH1753 | PRP8_Q1594A(TRP) FYV6_del ACT1-CUP1 A302C(3'ssUCG) | This work |
| yAAH3970 | yAAH3973 + pAAH1754 | PRP8_Q1594A(TRP) FYV6_del ACT1-CUP1 G303U(3'ssUAU) | This work |
| yAAH3971 | yAAH3973 + pAAH1755 | PRP8_Q1594A(TRP) FYV6_del ACT1-CUP1 G303C(3'ssUAC) | This work |
| yAAH3929 | ade2, cup1delta:ra3 his3 leu2 lys2 prp8delta:lys2 trp1 pJU169:PRP8(TRP) Fyv6::KAN | PRP8_WT(TRP) Prp18_V191A | This work |
| yAAH3930 | ade2, cup1delta:ra3 his3 leu2 lys2 prp8delta:lys2 trp1 pJU169:PRP8_Q1594A(TRP) Fyv6::KAN | PRP8_Q1594A(TRP) Prp18_V191A | This work |
| yAAH3948 | yAAH3929 + pAAH0470 | PRP8_WT(TRP) Prp18_V191A ACT1-CUP1 WT | This work |
| yAAH3949 | yAAH3929 + pAAH0526 | PRP8_WT(TRP) Prp18_V191A ACT1-CUP1 U301G(3'ssGAG) | This work |
| yAAH3950 | yAAH3929 + pAAH1644 | PRP8_WT(TRP) Prp18_V191A ACT1-CUP1 U301A(3'ssAAG) | This work |
| yAAH3951 | yAAH3929 + pAAH1645 | PRP8_WT(TRP) Prp18_V191A ACT1-CUP1 U301C(3'ssCAG) | This work |

| | | | |
|---|---|---|---|
| yAAH3952 | yAAH3929 + pAAH1752 | PRP8_WT(TRP) Prp18_V191A ACT1-CUP1 A302U(3'ssUUG) | This work |
| yAAH3953 | yAAH3929 + pAAH1753 | PRP8_WT(TRP) Prp18_V191A ACT1-CUP1 A302C(3'ssUCG) | This work |
| yAAH3954 | yAAH3929 + pAAH1754 | PRP8_WT(TRP) Prp18_V191A ACT1-CUP1 G303U(3'ssUAU) | This work |
| yAAH3955 | yAAH3929 + pAAH1755 | PRP8_WT(TRP) Prp18_V191A ACT1-CUP1 G303C(3'ssUAC) | This work |
| yAAH3940 | yAAH3930 + pAAH0470 | PRP8_Q1594A(TRP) Prp18_V191A ACT1-CUP1 WT | This work |
| yAAH3941 | yAAH3930 + pAAH0526 | PRP8_Q1594A(TRP) Prp18_V191A ACT1-CUP1 U301G(3'ssGAG) | This work |
| yAAH3942 | yAAH3930 + pAAH1644 | PRP8_Q1594A(TRP) Prp18_V191A ACT1-CUP1 U301A(3'ssAAG) | This work |
| yAAH3943 | yAAH3930 + pAAH1645 | PRP8_Q1594A(TRP) Prp18_V191A ACT1-CUP1 U301C(3'ssCAG) | This work |
| yAAH3944 | yAAH3930 + pAAH1752 | PRP8_Q1594A(TRP) Prp18_V191A ACT1-CUP1 A302U(3'ssUUG) | This work |
| yAAH3945 | yAAH3930 + pAAH1753 | PRP8_Q1594A(TRP) Prp18_V191A ACT1-CUP1 A302C(3'ssUCG) | This work |
| yAAH3946 | yAAH3930 + pAAH1754 | PRP8_Q1594A(TRP) Prp18_V191A ACT1-CUP1 G303U(3'ssUAU) | This work |
| yAAH3947 | yAAH3930 + pAAH1755 | PRP8_Q1594A(TRP) Prp18_V191A ACT1-CUP1 G303C(3'ssUAC) | This work |
| yAAH3933 | ade2, cup1delta:ura3 his3 leu2 lys2 prp8delta:lys2 trp1 pJU169:PRP8_WT(TRP) Prp22::HygR WT Prp22 (URA3/CEN) | PRP8_WT(TRP)WT Prp22 (URA/CEN) | This work |
| yAAH3935 | ade2, cup1delta:ura3 his3 leu2 lys2 prp8delta:lys2 trp1 pJU169:PRP8_Q1594A(TRP) Prp22::HygR WT Prp22 (URA3/CEN) | PRP8_Q1594A(TRP)WT Prp22 (URA/CEN) | This work |

| | | | |
|---|---|---|---|
| yAAH3974 | ade2, cup1delta:ura3 his3 leu2 lys2 prp8delta:lys2 trp1 pJU169:PRP8_WT(TRP) Prp22::HygR WT Prp22 (HIS3/CEN) | PRP8_WT(TRP)WT Prp22 (HIS/CEN) | This work |
| yAAH3975 | ade2, cup1delta:ura3 his3 leu2 lys2 prp8delta:lys2 trp1 pJU169:PRP8_Q1594A(TRP) Prp22::HygR WT Prp22 (HIS3/CEN) | PRP8_Q1594A(TRP)WT Prp22 (HIS/CEN) | This work |
| yAAH3989 | yAAH 3933 + pAAH1793 | PRP8_WT(TRP)WT Prp22 (URA/CEN) Prp22 S635A (HIS/CEN) | This work |
| yAAH3992 | yAAH 3933 + pAAH1796 | PRP8_WT(TRP)WT Prp22 (URA/CEN) Prp22 G810A (HIS/CEN) | This work |
| yAAH3993 | yAAH 3933 + pAAH1797 | PRP8_WT(TRP)WT Prp22 (URA/CEN) Prp22 I1133R (HIS/CEN) | This work |
| yAAH3987 | yAAH 3935 + pAAH1793 | PRP8_Q1594A(TRP)WT Prp22 (URA/CEN) Prp22 S635A (HIS/CEN) | This work |
| yAAH3994 | yAAH 3935 + pAAH1796 | PRP8_Q1594A(TRP)WT Prp22 (URA/CEN) Prp22 G810A (HIS/CEN) | This work |
| yAAH3995 | yAAH 3935 + pAAH1797 | PRP8_Q1594A(TRP)WT Prp22 (URA/CEN) Prp22 I1133R (HIS/CEN) | This work |
| yAAH4000 | yAAH 3933 + pAAH1799 | PRP8_WT(TRP)WT Prp22 (URA/CEN) Prp22 G810A I1133R (HIS/CEN) | This work |
| yAAH4001 | yAAH 3935 + pAAH1799 | PRP8_Q1594A(TRP)WT Prp22 (URA/CEN) Prp22 G810A I1133R (HIS/CEN) | This work |

**Supplementary Table S2.** Plasmids used in this study

| Stock ID | Plasmid name | Reference |
|---|---|---|
| pAAH1440 | pRS424 Prp8 WT TRP1 | This work |
| pAAH1626 | pRS424 Prp8_Q1594N TRP1 | This work |
| pAAH1627 | pRS424 Prp8_Q1594A TRP1 | This work |
| pAAH1628 | pRS424 Prp8_Q1594R TRP1 | This work |
| pAAH1629 | pRS424 Prp8_Q1594E TRP1 | This work |
| pAAH1630 | pRS424 Prp8_Q1594G TRP1 | This work |
| pAAH0470 | pGAC24-ACT1-CUP1 wild type | (Steinmetz and Brow 2003); gift from Dave Brow |
| pAAH1644 | pGAC24-ACT1-CUP1 U301A(3'ssAUG) | This work |
| pAAH1645 | pGAC24-ACT1-CUP1 U301C(3'ssCUG) | This work |
| pAAH1684 | pGAC24-ACT1-CUP1 U301G(3'ssGUG) | This work |
| pAAH1729 | pGAC24-ACT1-CUP1 3'ssGAGUAG | This work |
| pAAH1752 | pGAC24-ACT1-CUP1 A302U(3'ssUUG) | This work |
| pAAH1753 | pGAC24-ACT1-CUP1 A302C(3'ssUCG) | This work |
| pAAH1754 | pGAC24-ACT1-CUP1 G303U(3'ssUAU) | This work |
| pAAH1755 | pGAC24-ACT1-CUP1 G303C(3'ssUAC) | This work |
| pAAH1761 | pGAC24-ACT1-CUP1 3'ssUAGUAG | This work |
| pAAH1762 | pGAC24-ACT1-CUP1 3'ssUAGGAG | This work |
| pAAH0278 | pAG25-NATMX | (Goldstein and McCusker 1999); Euroscarf |
| pAAH0093 | pUG6-KANMX | (Goldstein and McCusker 1999); Euroscarf |
| pAAH1603 | p360-Prp18-11(Prp18V191A) | (Aronova et al. 2007); gift from Beate Schwer |
| pAAH1791 | pRS314-WT Prp22 His | This work |
| pAAH1793 | pRS314-Prp22 S635A His | This work |
| pAAH1796 | pRS314-Prp22 G810A His | This work |
| pAAH1797 | pRS314-Prp22 I1133R His | This work |
| pAAH1799 | pRS314-Prp22_G810A I1133R His | This work |
